# Membrane potential bistability in human breast cancer MDA-MB-231 cells: A ‘Hodgkin-Huxley type’ model

**DOI:** 10.1101/2025.07.03.663062

**Authors:** Suvabrata De, Mustafa B. A. Djamgoz

## Abstract

The plasma membrane voltage (*V*_m_) is well known to have significant involvement in a wide range of cellular functions including cancer progression. Voltage imaging revealed that *V*_m_ of MDA-MB-231 breast cancer cells is ‘bistable’ with hyperpolarising voltage transients (HVTs). Here, we formulate a model of *V*_m_ incorporating the ion channels Na_v_1.5, Ca_v_3.2, and K_Ca_1.1. *V*_m_ is governed by the Hodgkin-Huxley formalism coupled to intracellular Ca^2+^ dynamics, via Ca^2+^ influx through Ca_v_3.2 and Ca^2+^-dependent efflux of K^+^ through K_Ca_1.1. Stochastic fluctuations—arising from sparse ion channel expression and Ca^2+^-induced Ca^2+^ release (CICR)—drive *V*_m_ transitions between the otherwise stable depolarised and hyperpolarised states. The model qualitatively reproduces the key experimental observations of HVTs, and their suppression by specific inhibitors of Na_v_1.5 or K_Ca_1.1. It is predicted that inhibition of CICR should also lead to suppression of HVTs. Our model promises to help the understanding of the dynamic electrical activity of the MDA-MB-231 cell model and its functional consequences, and may inspire future bioelectricity-based cancer diagnosis and therapy.

## 1 Introduction

Several studies have shown that ion channels play a significant role in cancer cell behaviour, including proliferation and invasion (Fraser et al. 2005; Levin 2021; Prevarskaya, Skryma, and Shuba 2018). In particular, cancer cells with the ability to metastasise have electrically active membranes driven in part by de novo expression of voltage-gated sodium channels (VGSCs) (Djamgoz 2024).

Most work has been done on breast cancer—especially the MDA-MB-231 cell line model—where the dominant VGSC is the neonatal splice variant of Na_v_1.5 (Fraser et al. 2005). Thus, current clamp and multi-electrode array recordings have shown that these cells display dynamic electrical activity, which is blocked reversibly by tetrodotoxin (TTX) (Ribeiro et al. 2020). Most recently, the membrane potential (*V*_m_) of MDA-MB-231 cells has been studied by voltage imaging, which showed that the cells undergo hyperpolarising voltage transients (HVTs)—spontaneous ‘negative going events’ of *V*_m_—suggesting *V*_m_ bistability: the existence of hyperpolarised and depolarised resting *V*_m_s (Quicke et al. 2022). Pharmacological intervention revealed that the HVTs involve VGSC and voltage- and calcium-(Ca^2+^) activated potassium (K^+^) channel (K_Ca_1.1) activities (Quicke et al. 2022).

In this paper, we develop and analyse a minimal model for bistability in MDA-MB-231 cells, wherein *V*_m_ evolves according to the Hodgkin-Huxley (HH) formalism (Hodgkin and Huxley 1952) and is coupled to intracellular Ca^2+^ dynamics. The essential ion channels in our model are Na_v_1.5, Ca_v_3.2 and K_Ca_1.1; the existence of inward Ca^2+^ current (through Ca_v_3.2) and the dependence of outward K^+^ current (through K_Ca_1.1) on Ca^2+^ results in coupling between *V*_m_ and Ca^2+^. Additionally, the dynamical equations are subject to stochastic fluctuations, which induce transitions between hyperpolarised and depolarised resting *V*_m_s. Fluctuations in ion channel dynamics (gating variables) arise due to the relatively sparse expression (low density) of the ion channels, in particular K_Ca_1.1, while the phenomenon of Ca^2+^-induced Ca^2+^ release (CICR) (Fabiato 1983; Endo 2009) motivates the inclusion of multiplicative noise in Ca^2+^ dynamics. We find that the predicted dynamics of *V*_m_ successfully capture the main observations of voltage imaging: the occurrence of HVTs, and the disappearance of bistability and HVTs upon the pharmacological blocking of Na_v_1.5 and K_Ca_1.1 (Quicke et al. 2022). Furthermore, the model yields the novel prediction that inhibition of CICR would induce hyperpolarisation, and thus suppress HVTs.

A number of theoretical studies have investigated the role of bioelectric signalling and spatio-temporal patterning in multicellular cancer biophysics (Cervera, Alcaraz, and Mafe 2016; Carvalho 2021; Cervera, Levin, and Mafe 2023). In the context of breast cancer, prior work has focused on the functional importance of individual ion channels or aspects of Ca^2+^ signalling, and we are unaware of any studies—either experimental or theoretical—that have analysed the integrated effect of several ion channels and intracellular Ca^2+^ dynamics. Thus, to the best of our knowledge, the current work represents the first mathematical model of the dynamic behaviour of *V*_m_ and Ca^2+^ within a breast cancer cell line.

## 2 The model

We now specify the elements that comprise our minimal model. In Section 2.1, we outline the gating regulation and dynamics of our chosen ion channels, and how they govern the evolution of *V*_m_ via the HH formalism. Section 2.2 deals with the dynamics of intracellular Ca^2+^, which couples to the dynamics of *V*_m_.

### 2.1 V_m_ dynamics

#### 2.1.1 Na_v_1.5

The connection between upregulated functional VGSC expression and electrical activity in cancer cells has been argued for over two decades (Fraser et al. 2005; Djamgoz 2024). MDA-MB-231 cells have been shown to express the VGSC Na_v_1.5, in the form of an embryonic splice variant (‘neonatal’ Na_v_1.5) (Fraser et al. 2005). The subtle electrophysiological differences between the adult form of Na_v_1.5—the cardiac isoform—and the neonatal form have been investigated (Onkal et al. 2008). Models of cardiac electrical activity have successfully utilised ‘*m*^3^*h*’ dynamics (introduced by Hodgkin and Huxley) for adult Na_v_1.5 (Di Francesco and Noble 1985). In light of this, we assume that modelling neonatal Na_v_1.5 in this way is valid for our purposes as well ^1^.

The activation (*m*_Na_) and inactivation (*h*_Na_) gating variables of Na_v_1.5 determine the dynamical changes in conductance of the channel. Roger et al. measured the steady-state (equilibrium) forms of the gating variables to be as follows:

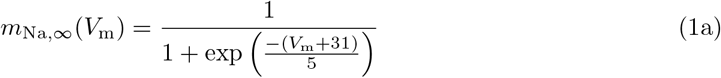

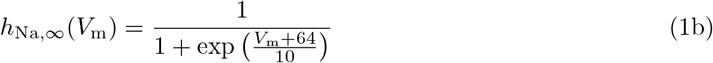

and these are displayed in Fig. 1a (Roger, Besson, and Le Guennec 2003). The constants 5 mV and 10 mV are slope factors which determine the steepness of the curves, and were estimated by eye from the measured data (Roger, Besson, and Le Guennec 2003). In general, the gating variables will not be at equilibrium, and so to determine their dynamics, the *V*_m_-dependent time constants of (in)activation 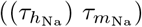 are required. The work of Zhang et al. in modelling the kinetics of adult Na_v_1.5 (Zhang et al. 2013) enables us to approximate these as follows (in ms ^2^):

**Figure 1:**
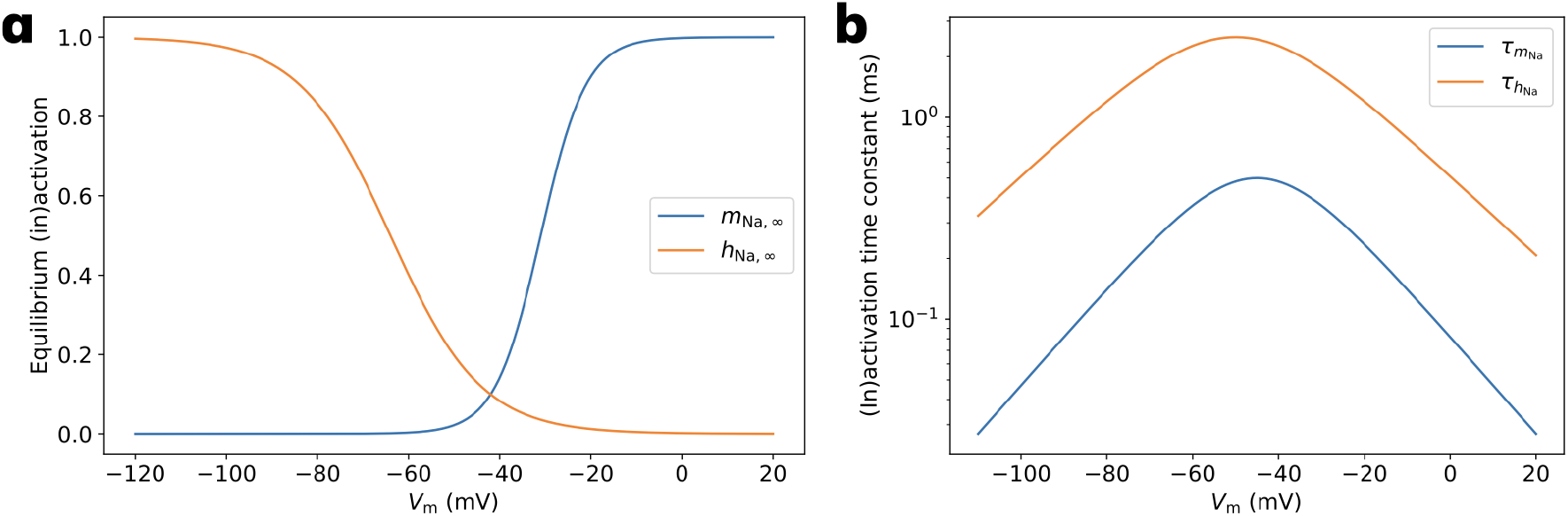
Gating variables and time constants for Na_v_1.5. **(a)** Steady state (in)activation ((*h*_Na,*∞*_) *m*_Na,*∞*_) gating variables (Eq. 1) as measured by Roger et al. (Roger, Besson, and Le Guennec 2003). Note the gradual decrease in *h*_Na,*∞*_ for depolarised *V*_m_, which enables Na_v_1.5 to be functionally active at ∼ −20 mV. **(b)** *V*_m_-dependent time constants of (in)activation 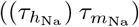—see Eq. 2—estimated from (Zhang et al. 2013).

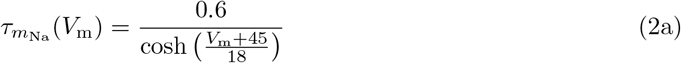

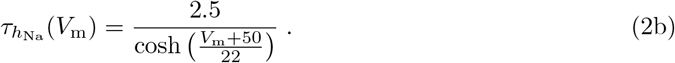

This is in line with the functional form commonly used in biophysical models of *V*_m_ dynamics (G. C. Smith 2019) ^3^; the *V*_m_-dependency of the time constants is shown in Fig. 1b. Thus, *m*_Na_ and *h*_Na_ obey the following ordinary differential equations (ODEs):

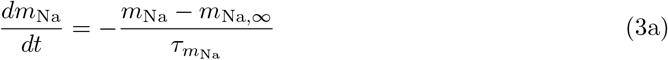

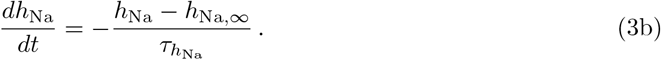

The sodium current density ^4^ is thus given by

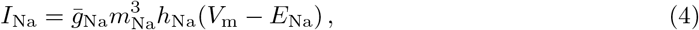

where 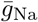 and *E*_Na_ are the maximum conductance density and sodium Nernst potential, respectively, and their values are shown in Table 1. Using Eq. 4, we estimate 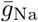 from (i) the data of Fraser et al. (inset of their Fig. 1B) (Fraser et al. 2005); (ii) the values of *E*_Na_ and cell radius (*r*) (Table 1); (iii) the assumption that at the peak of the control current trace, *m*_Na_ ≃ 1 and *h*_Na_ ≃ 1.

**Table 1:** Values of parameters used in the model.

| Parameter | Value | Reference(s) |
| --- | --- | --- |
| $\bar{g}_{Na}$ | 0.85 mS/cm<sup>2</sup> | (Fraser et al. 2005) |
| $\bar{g}_{Ca}$ | 0.052 mS/cm<sup>2</sup> | (Taylor et al. 2008) |
| $\bar{g}_K$ | 1.1 mS/cm<sup>2</sup> | (Sizemore et al. 2020) |
| $N_{Na}$ | 537 | (Kang et al. 2021; Selimi et al. 2023) |
| $N_{Ca}$ | 368 | (Engbers, Zamponi, and Turner 2013) |
| $N_K$ | 71 | (Contreras et al. 2013) |
| $E_{Na}$ | 55 mV | |
| $E_K$ | -75 mV | |
| $c_o$ | 1 mM | |
| $c_{rest}$ | $1.5 \cdot 10^{-5}$ mM | (Rüdiger, Shuai, and Sokolov 2010; Dobramysl, Sten Rüdiger, and Erban 2016) |
| $\gamma$ | $10^{-4}$ ms<sup>-1</sup> | (Pieri et al. 2013; Wolf and Wennemuth 2014) |
| $f$ | 0.005 | (Chay and Keizer 1983; Keener and Sneyd 2008) |
| $C$ | 1.6 μF/cm<sup>2</sup> | (Han, L. Yang, and Frazier 2007; Awad et al. 2024) |
| $r$ | $10^{-3}$ cm | (Morley, Walsh, and Newport 2017) |
| $F$ | $9.6485 \cdot 10^4$ C/mol | |
| $R$ | 8.3145 J/(mol K) | |
| $T$ | 295.15 K | |

#### 2.1.2 Ca_v_3.2

The physiological importance of voltage-gated Ca^2+^ channels (VGCCs) in various excitable cells—including neurons, cardiac and endocrine cells—is well appreciated, as reviewed in (Catterall 2011). Moreover, within the last two decades, evidence has accumulated suggesting the pathophysiological role of such channels in cancer cells, particularly with respect to proliferation (Prevarskaya, Skryma, and Shuba 2018; Bhargava and Saha 2019). The T-type VGCC Ca_v_3.2 was first demonstrated to be significantly expressed in MDA-MB-231 cells by Gray et al. in 2004 (Gray et al. 2004). Taylor et al. replicated this finding and additionally showed that selectively blocking Ca_v_3.2 suppresses proliferation (Taylor et al. 2008).

We follow Engbers et al. (Engbers, Zamponi, and Turner 2013) in assuming that the channel obeys ‘*mh*’ dynamics, with equilibrium (in)activation ((*h*_Ca_) *m*_Ca_) gating variables given by

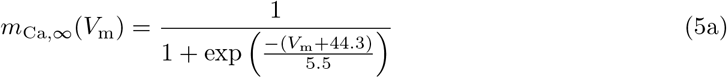

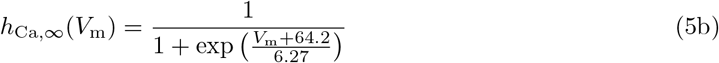

as displayed in Fig. 2. The time constants of (in)activation 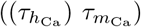 presented in (Engbers, Zamponi, and Turner 2013) are

**Figure 2:**
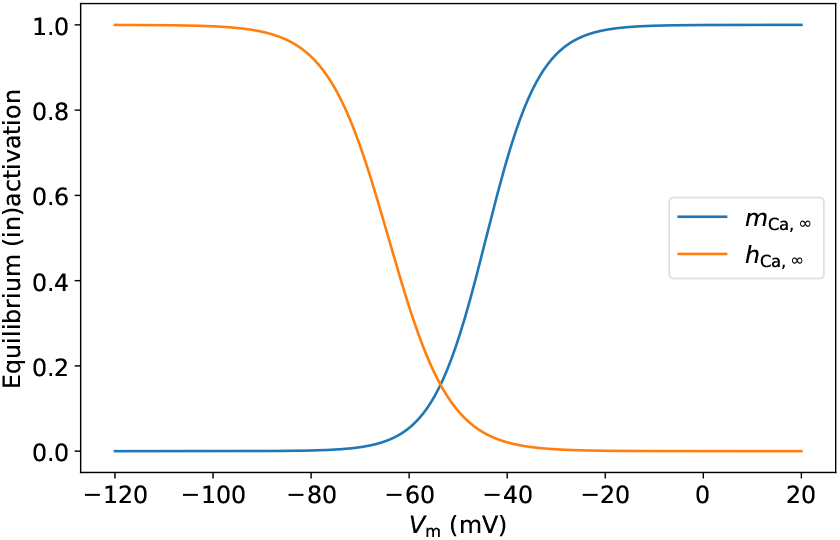
Gating variables for Ca_v_3.2. Steady state (in)activation ((*h*_Ca,*∞*_) *m*_Ca,*∞*_) gating variables (Eq. 5) as utilised by Engbers et al. (Engbers, Zamponi, and Turner 2013). Note that Ca_v_3.2 channels remain functionally active at low (∼ −50 mV) values of *V*_m_.

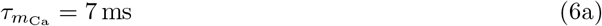

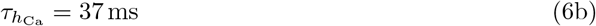

enabling us to write the following ODEs:

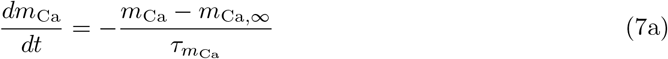

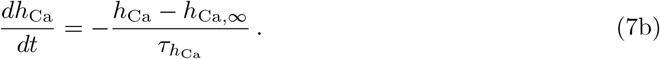

We thus express the Ca^2+^ current as

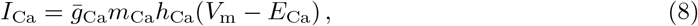

where 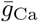 is the maximum conductance density (Table 1), and the Ca^2+^ Nernst potential varies with changes in the intracellular Ca^2+^ concentration (*c*_i_) as follows ^5^

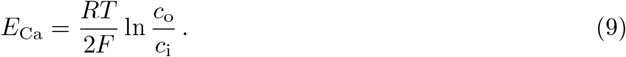

In Eq. 9, the values of *R* (the gas constant), *T* (absolute temperature) ^6^, *F* (Faraday’s constant) and *c*_o_ (the fixed extracellular Ca^2+^ concentration) are presented in Table 1. We estimate 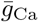 using Eq. 8 with the help of: (i) the data from Fig. 4A in (Taylor et al. 2008); (ii) the experimental condition of *c*_o_ = 5 mM (Taylor et al. 2008) ^7^; (iii) the assumption of *c*_i_ = 10^*−*4^ mM (yielding *E*_Ca_ = 138 mV); (iv) the assumption that *m*_Ca_ ≃ 1 and *h*_Ca_ ≃ 1 at the peak of the control current trace displayed in Fig. 4A from (Taylor et al. 2008).

#### 2.1.3 K_Ca_1.1

The large-conductance voltage- and Ca^2+^-activated K^+^ channel K_Ca_1.1 (also known as BK) is widely expressed in excitable and non-excitable cells of mammalian tissues (Contreras et al. 2013). Studies have shown correlation between over-expression of *KCNMA1* (encoding the *α*-subunit of K_Ca_1.1) and the pathophysiology of breast cancers, including metastasis and invasion (Khaitan et al. 2009) and proliferation (Oeggerli et al. 2012) ^8^. Indeed, it is well established that K_Ca_1.1 is highly expressed in MDA-MB-231 cells (Roger, Potier, et al. 2004; Khaitan et al. 2009; M. Yang et al. 2020; Sizemore et al. 2020). Interestingly, there is evidence that K_Ca_1.1 and Ca_v_3.2 co-localise (form complexes) in prostate cancer cells (Gackiere et al. 2013) as well as in rat medial vestibular neurons *in vitro* (Rehak et al. 2013). Sizemore et al. present evidence that the activity of K_Ca_1.1 channels dominates that of other K^+^ channels, by applying the specific K_Ca_1.1 blocker iberiotoxin (IbTX) (Sizemore et al. 2020). This justifies the inclusion of K_Ca_1.1 as the only hyperpolarising channel (permitting outward current) in our model.

We employ the relatively simple dynamics of K_Ca_1.1 proposed by Yamada et al. in their modeling of bullfrog sympathetic ganglion cells (Yamada, Koch, and Adams 1989). The channel is non-inactivating, and the steady-state activation (*m*_K,*∞*_) and activation time constant 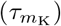 are *c*_i_- and *V*_m_-dependent. These are conveniently expressed in terms of the forward 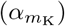 and backward 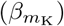 rate constants,^9^ given by

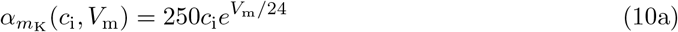

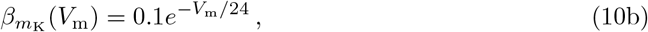

from which follows

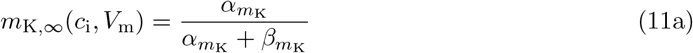

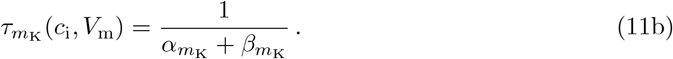

The surface plots of the steady-state activation and activation time constant are displayed in Fig. 3. *m*_K_ thus obeys the ODE

**Figure 3:**
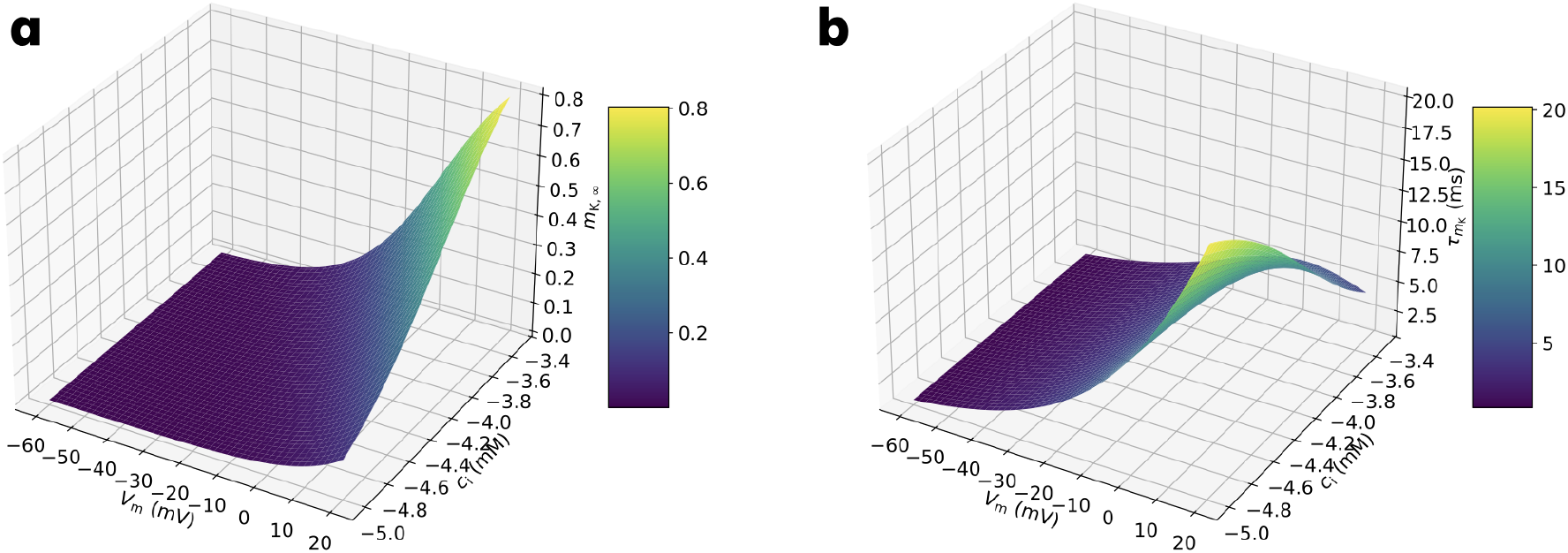
Activation variable and associated time constant for K_Ca_1.1. **(a)** Steady state activation (*m*_K,*∞*_) gating variable (Eq. 11a) as utilised by Yamada et al. (Yamada, Koch, and Adams 1989). Despite *m*_K,*∞*_ being small (*<* 10^*−*2^) for typical values of *c*_i_ and *V*_m_ in our model (e.g. *c*_i_ ∼ 10^*−*5^ mM and *V*_m_ ∼ −20 mV), K_Ca_1.1 channels remain functionally active due to their very large single-channel conductance (see Section 2.1.4). **(b)** The *c*_i_- and *V*_m_-dependent time constant of activation 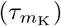—see Eq. 11b (Yamada, Koch, and Adams 1989). 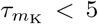 ms for typical values of *c*_i_ and *V*_m_ in our model.

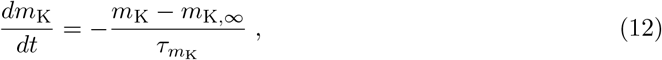

the solution of which allows the K_Ca_1.1 current to be computed as

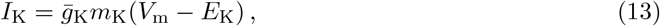

where 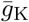 is the maximum conductance density and *E*_K_ is the K^+^ Nernst potential (Table 1). The work of Sizemore et al. (Fig. 3A in (Sizemore et al. 2020)) permits us to estimate 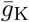 using Eq. 13: for the depolarising pulse of 50 mV, it can be assumed that after 300 ms, the activation variable reaches its steady-state value close to unity, *m*_K_ ≃ 1 ^10^.

#### 2.1.4 Deterministic and stochastic Hodgkin-Huxley formalism

With the help of Eqs. 4, 8 and 13, we arrive at the current-balance equation (analogous to that used in the original HH model (Hodgkin and Huxley 1952)):

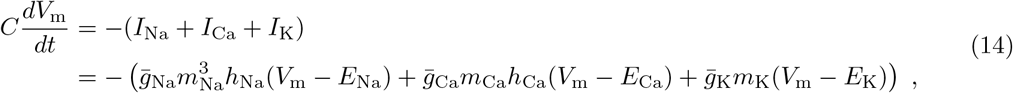

where *C* is the membrane capacitance (Table 1). Eq. 14 along with Eqs. 3, 7 and 12 are deterministic equations that comprise our HH-type modelling of *V*_m_ dynamics. However, as mentioned in Section 1, the low numbers of ion channels expressed in the membrane of MDA-MB-231 cells (compared to typical neurons) means that the channel conductances, and thus *V*_m_, are subject to stochastic fluctuations. Indeed, from our estimates of the maximum conductance densities, we can estimate the number of ion channels of each type (*N*_X_, X = Na, Ca or K; see Table 1), with the aid of data on single-channel (unitary) conductances of adult Na_v_1.5 (20 pS ^11^), Ca_v_3.2 (1.7 pS ^12^) and K_Ca_1.1 (200 pS ^13^).

Conductance fluctuations due to relatively low numbers of channels have been modelled via Markov processes, wherein channels transition in an independent and random fashion between distinct configurations (Groff, DeRemigio, and G. D. Smith 2009). Fox and Lu derived stochastic differential equations (SDEs) which approximate such Markov process models (Fox and Lu 1994), and have the desirable properties of being simpler and computationally more efficient. Specifically, they showed that channel gating ODEs (such as Eqs. 3, 7 and 12) can be replaced by a SDE of the form

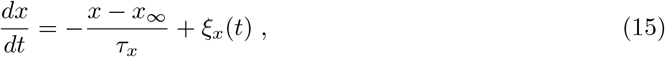

where *x* = *m*_Na_, *h*_Na_, *m*_Ca_, *h*_Ca_ or *m*_K_; *ξ*_*x*_(*t*) represents statistically independent Gaussian white noise, with zero mean and second moment given by

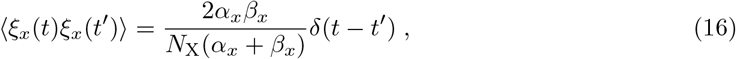

where *δ*(*t* − *t*^*′*^) is the Dirac delta function, while *α*_*x*_ and *β*_*x*_ can be expressed in terms of *x*_*∞*_ and *τ*_*x*_ from relations analogous to Eq. 11 for each gating variable *x*. Now that the gating variables in Eq. 14 are understood to be stochastic variables, *V*_m_ evolves stochastically as well. Eqs. 14 and 15 define the stochastic HH formalism we employ in our modelling.

### 2.2 Ca^2+^ dynamics

#### 2.2.1 Deterministic dynamics

At present, relatively little is known about the mechanistic details of intracellular Ca^2+^ dynamics in MDA-MB-231 cells (Rizaner et al. 2016). In light of this, we model the deterministic dynamics of free (unbuffered) cytosolic Ca^2+^ by accounting for influx due to current (*I*_Ca_) through Ca_v_3.2 channels, and for vanishingly small current, a tendency for *c*_i_ to decay to a physiological resting concentration, *c*_rest_ (Table 1) ^14^. The latter feature minimally captures the regulatory effect of pumps, exchangers and other transport mechanisms in maintaining *c*_i_ ≃ *c*_rest_. We thus utilise the following ODE:

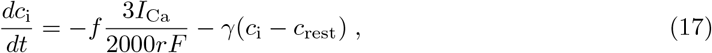

where *γ* is the decay (clearance) rate of Ca^2+^ from the cytoplasm, and *f* is a buffering scale factor, reflecting the binding of the majority (∼ 99%) of Ca^2+^ entering the cell to cytoplasmic proteins (see Table 1 for the values of *γ* and *f* we employ) ^15^. Note that in Eq. 17, *I*_Ca_ is time- and *V*_m_-dependent, thus coupling the dynamics of Ca^2+^ to *V*_m_, while a negative current (influx of Ca^2+^) corresponds to a positive change in *c*_i_. Our modelling of the deterministic dynamics of Ca^2+^ is similar to that employed by Chay and Keizer in their model of *V*_m_ oscillations in pancreatic *β*-cells (Chay and Keizer 1983).

#### 2.2.2 CICR and stochastic dynamics

It is well known that a key regulatory mechanism controlling cytosolic Ca^2+^ is the release of Ca^2+^ from internal stores such as the endoplasmic reticulum (ER) (Dupont, Falcke, et al. 2016). This is facilitated by two families of channels, namely, Ryanodine receptors (RyRs) and Inositol (1,4,5)-trisphosphate receptors (IP_3_Rs). A key characteristic of both receptors is that their probability of being open is modulated by Ca^2+^ itself. Thus, a small increase in Ca^2+^ concentration (beyond a threshold) at the mouth of these channels can trigger a fast positive feedback process resulting in substantial Ca^2+^ release; this phenomenon (alluded to in Section 1) is known as CICR. Clearly however, the positive feedback process cannot continue indefinitely: for IP_3_Rs, a slower negative feedback process wherein Ca^2+^ inactivates the channel ensues, terminating the release of Ca^2+^. A combination of the buffering of Ca^2+^, its removal by pumps, and Ca^2+^ diffusion throughout the cytoplasm, thus reduces the Ca^2+^ concentration over a longer timescale. In this way, the mechanism described forms an excitable system giving rise to Ca^2+^ transients or spikes—analogous to neuronal action potentials (APs)—and has inspired models of such behaviour utilising the HH formalism (Li and Rinzel 1994).

In their study of breast cancer specimens, Abdul et al. found a correlation between RyR expression and tumour grade, and furthermore, demonstrated *in vitro* that the RyR agonist 4-chloro-*m*-cresol inhibited proliferation of the cell lines MDA-MB-231 and MCF-7 (Abdul, Ramlal, and Hoosein 2008). Evidence of the significance of type 3 IP_3_R (IP_3_R3) in breast cancers has accumulated. In the MCF-7 cell line, inhibition of IP_3_R3 was shown to reduce proliferation (Szatkowski et al. 2010), while a follow-up study implicated IP_3_R3-K_Ca_1.1 co-localisation and interaction with increased proliferation (Mound, Rodat-Despoix, et al. 2013). Subsequently, the silencing of IP_3_R3 has been shown to significantly reduce the migration of various cell lines, including MDA-MB-231 (Mound, Vautrin-Glabik, et al. 2017).

We incorporate CICR in the dynamics of Ca^2+^ phenomenologically, with the addition of a multiplicative noise term to Eq. 17 (thus rendering *c*_i_ a stochastic variable) as follows:

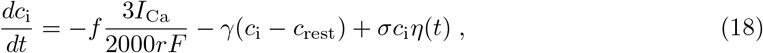

where *η*(*t*) is statistically independent Gaussian white noise, with zero mean and an autocorrelation of ⟨*η*(*t*)*η*(*t*^*′*^)⟩ = *δ*(*t* − *t*^*′*^). Furthermore, in Eq. 18, the constant *σ* sets the strength of the noise, while the noise amplitude scales with *c*_i_, reflecting the positive feedback nature of CICR.

## 3 Results

### 3.1 Bistability

To determine the steady state (equilibrium) values of *V*_m_ and *c*_i_ predicted by our model, we examine their deterministic evolution, Eqs. 14 and 17. The inclusion of stochasticity (via the corresponding stochastic equations) will be seen to induce local fluctuations around the steady states, or, promote transitions between them. Steady state solutions of Eqs. 14 and 17 are values of *V*_m_ and 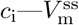 and 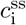, respectively—for which 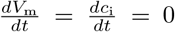 indefinitely, and not instantaneously ^16^. 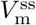 and 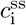 are found numerically by solving both equations simultaneously after replacing the gating variables with their steady state expressions (e.g. *m*_Na_ → *m*_Na,*∞*_). 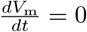 defines a curve in the *c*_i_-*V*_m_ plane, while 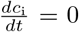 defines another curve ^17^; the intersection(s) of these curves yields the solution(s) 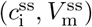, as shown in Fig. 4. Three points of equilibrium are apparent: a depolarised state with *c*_i_ close to *c*_rest_, (2.1 · 10^*−*5^ mM, −20 mV), a hyperpolarised state at a ‘high’ *c*_i_, (1.7 · 10^*−*4^ mM, −49 mV), and an intermediate state, (4.1 · 10^*−*5^ mM, −30 mV).

**Figure 4:**
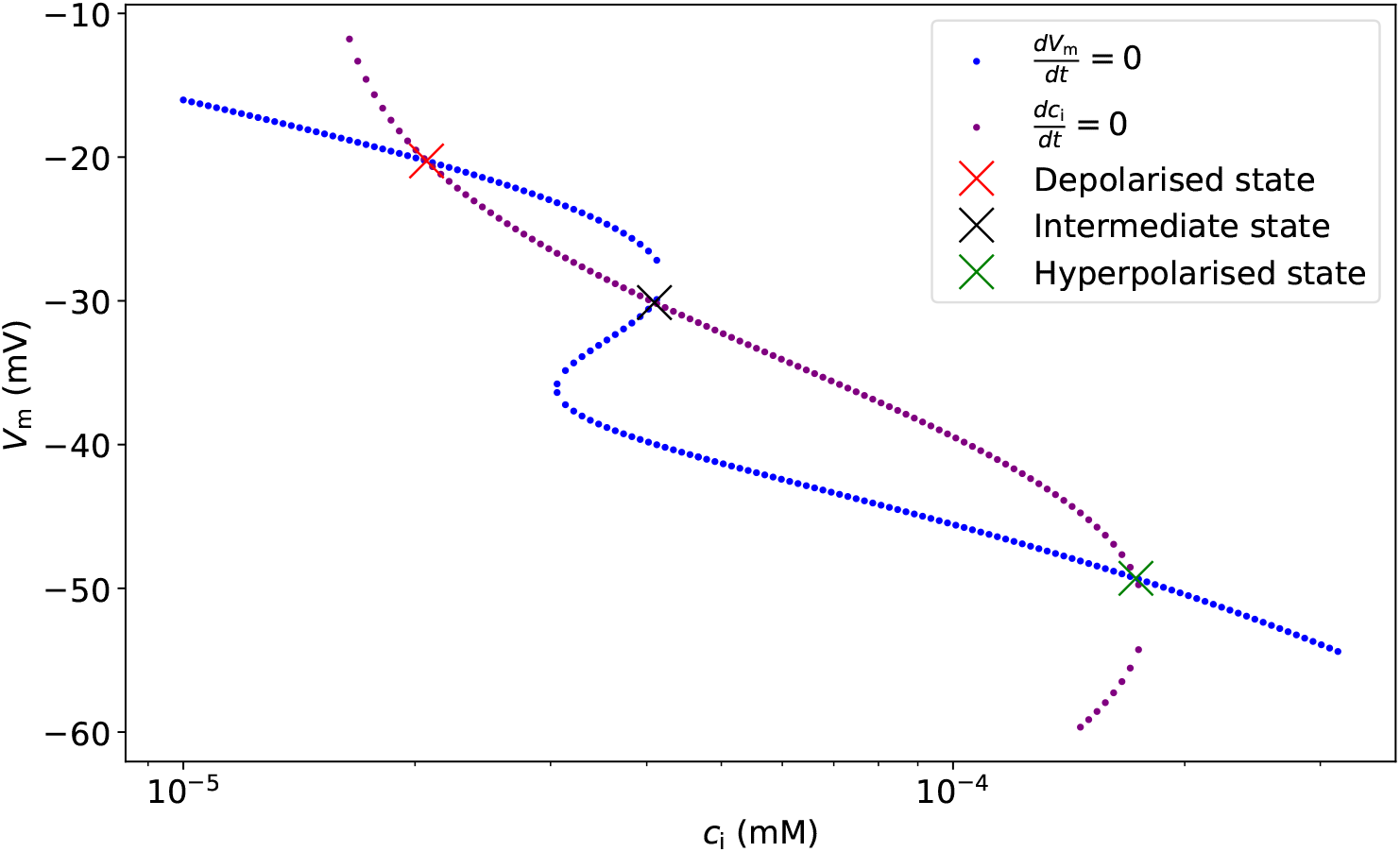
Nullclines for *c*_i_ (purple dots) and *V*_m_ (blue dots). The three points of intersection of the nullclines (marked by crosses) correspond to the three steady states of the system.

The nature of these equilibria can be understood with the aid of phase planes: visualisations of the system’s dynamics (Strogatz 2018). Fig. 5 displays phase planes in the neighbourhood of the depolarised and intermediate steady states ^18^. At each point in the *c*_i_-*V*_m_ planes, the corresponding vector depicts the local rates of change of *c*_i_ and *V*_m_; longer vectors represent greater rates of change, while a non-existent vector represents equilibrium at that point in the plane. Fig. 5a shows that the system evolves (‘flows’) toward equilibrium, whereas the flows in Fig. 5b are away from equilibrium (except for points that fall on an almost horizontal line passing through equilibrium). Local perturbations away from the depolarised steady state (Fig. 5a) thus return to equilibrium: the steady state is stable, as is the hyperpolarised equilibrium. On the other hand, generic local perturbations away from the intermediate steady state (Fig. 5b) do not: this is an unstable state, and thus, the system is bistable.

**Figure 5:**
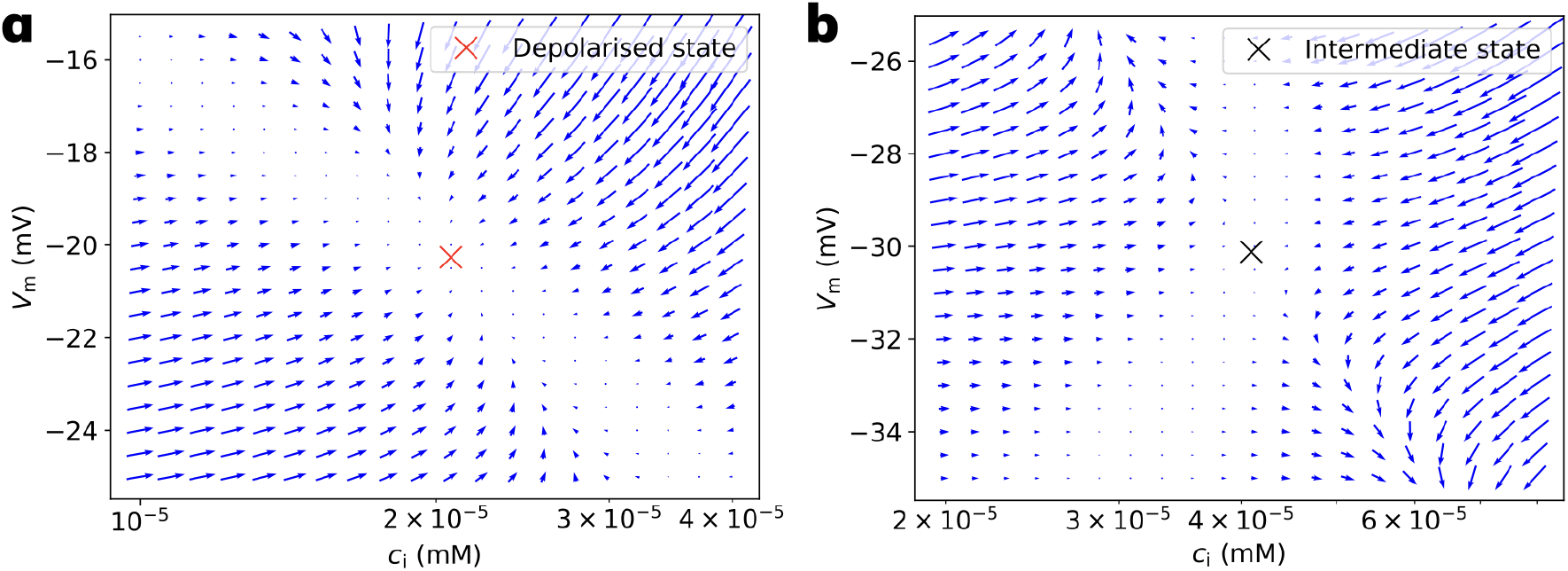
Phase planes in the vicinity of the depolarised (a) and intermediate (b) steady states. Points of equilibrium are marked by crosses. For clarity, 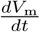 is scaled by 10^*−*3^, which does not alter the nature of stability of either state.

### 3.2 HVTs

We now consider the stochastic dynamics of *V*_m_ and *c*_i_, by integrating the system of SDEs (Eqs. 14, 15 and 18) with the Euler-Maruyama method and a stepsize of Δ*t* = 10^*−*3^ ms (Higham and Kloeden 2021). The parameters used are those in Table 1, while *σ* is the only adjustable parameter of our model, which will be seen to affect the qualitative nature of the dynamics. The initial conditions of all simulations are the depolarised state (e.g.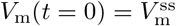) with the gating variables set to their equilibrium values (e.g.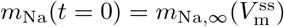).

An example of a 100 s simulation of the system of SDEs for *σ* = 0.05 ms^*−*1*/*2^ is shown in Fig. 6. Examination of the *V*_m_- and *c*_i_-time series reveals that typically, the system fluctuates in a state of depolarisation with correspondingly low *c*_i_. For example, during the interval *t* = 70 s - 90 s, the average state variables are 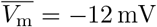 and 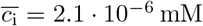. At such depolarised values of *V*_m_, Ca_v_3.2 channels are almost completely inactivated (Fig. 2) resulting in negligible *I*_Ca_. In contrast, Na_v_1.5 and K_Ca_1.1 channels support finite (albeit fluctuating) currents (Fig. 6c), which on average counterbalance to maintain a noisy depolarised equilibrium. Such intervals correspond to noisy excursions in the vicinity of the depolarised steady state in the *c*_i_-*V*_m_ plane (Fig. 4).

**Figure 6:**
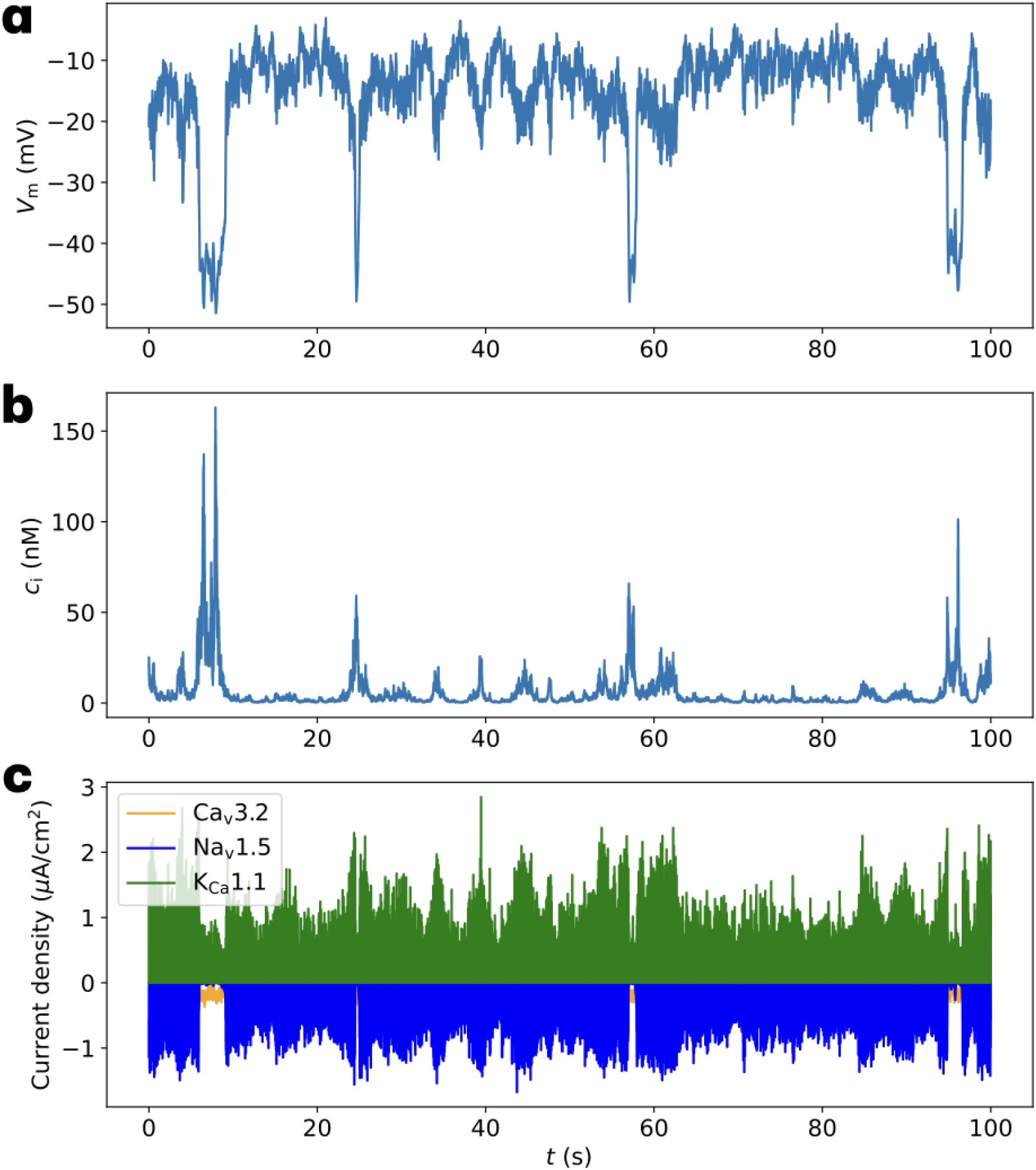
A 100 s simulation of the SDEs displaying HVTs. Stochastic version of Eq. 14 and Eq. 18 simulated for *σ* = 0.05 ms^*−*1*/*2^, with corresponding time series for *V*_m_ **(a)**, *c*_i_ **(b)** and currents for each ionic species **(c)**. Four HVTs (e.g. at ∼ 25 s) can be seen in **(a)**, with synchronous spikes of *c*_i_ seen in **(b). (c)** demonstrates that HVTs coincide with a reduction in *I*_Na_ to negligible values, and a corresponding increase in *I*_Ca_.

Fig. 6a displays that intervals of depolarisation are punctuated by relatively brief and sudden HVTs, where *V*_m_ hyperpolarises abruptly from ∼ −20 mV to ∼ −50 mV, and subsequently depolarises. The absolute change in *V*_m_ during a HVT— |Δ*V*_m_| ≃ 30 mV—is comparable to the changes recorded experimentally (Quicke et al. 2022). One can see that HVTs are accompanied by spikes (transients) in *c*_i_ (Fig. 6b) and dramatic changes in the current flows (Fig. 6c). Indeed, we can identify six key steps in a HVT (Fig. 7); examining these steps by ‘zooming in’ on the HVT between 24.4 s and 25.2 s (Fig. 8a-c), for example, is instructive.

**Figure 7:**
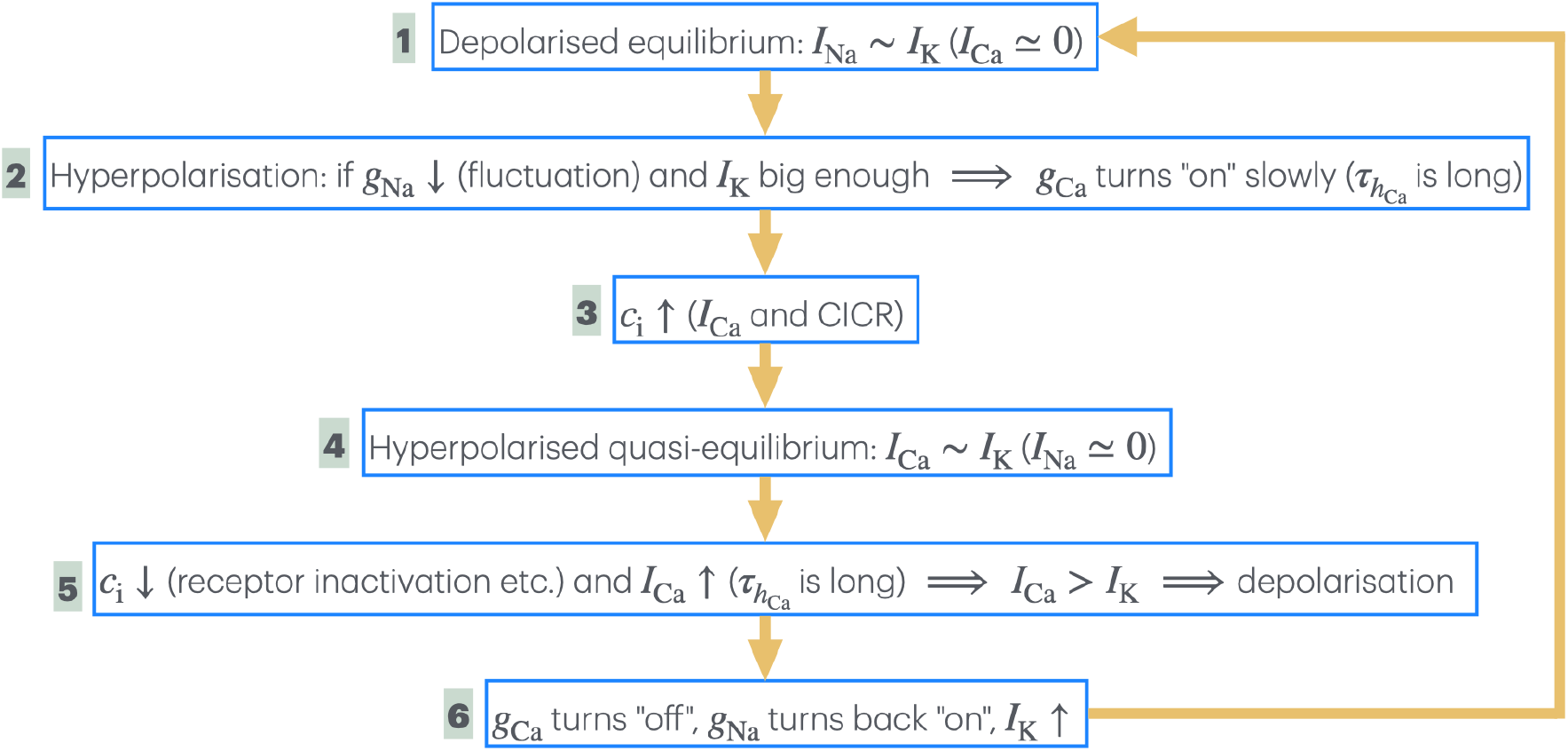
The six cyclical steps in a HVT. The distinct steps in the transition from depolarised equilibrium to hyperpolarised equilibrium (and vice-versa) are labelled 1 to 6. To avoid notational clutter, *I*_X_ refers to the magnitude of current of ionic species X. *g*_Na_ is the dynamical conductance density of Na_v_1.5 channels, i.e.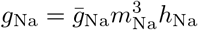, with *g*_Ca_ having a similar meaning. The arrow from step 6 to step 1 denotes repetition of the cyclical process, and should not be interpreted as feedback.

**Figure 8:**
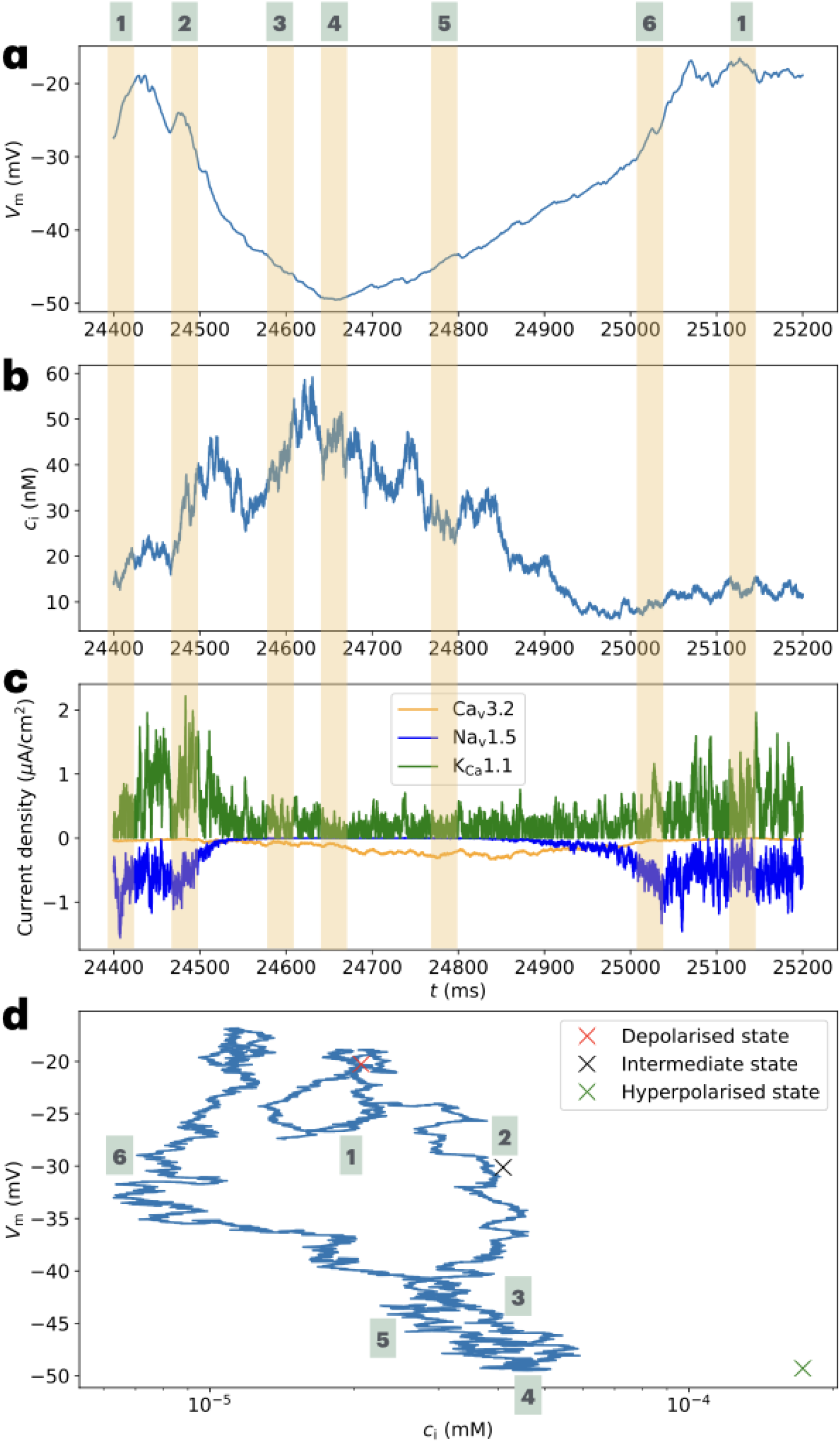
Temporal magnification and trajectory of a HVT. **(a), (b), (c)** A fragment of the simulation displayed in Fig. 6 between 24.4 s and 25.2 s. The six distinct steps of the HVT (with numbering at the top corresponding to Fig. 7) are highlighted within the time series for *V*_m_ **(a)**, *c*_i_ **(b)** and ionic currents **(c). (d)** The system’s trajectory in the *c*_i_-*V*_m_ plane between 24.4 s and 25.1 s of the simulation displayed in Fig. 6, with the steps of the HVT highlighted.

Prior to hyperpolarisation, the cell exists in a state of depolarised equilibrium, where *I*_Na_ counterbalances *I*_K_ on average, while *I*_Ca_ is negligible (step 1 - Fig. 8c). If a stochastic fluctuation causes an appreciable decrease in the conductance of Na_v_1.5, and simultaneously, *I*_K_ is large enough, substantial hyperpolarisation occurs (step 2 - Figs. 8a,c). This results in the conductance of Ca_v_3.2 increasing slowly, due to its long (de)inactivation time constant, 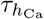 (Eq. 6b). Hyperpolarisation and the resulting increase in *I*_Ca_ leads to an influx of Ca^2+^, which catalyses CICR and results in a rapid rise of *c*_i_ (step 3 - Fig. 8b). Subsequently, a brief state of hyperpolarised equilibrium (‘quasi-equilibrium’) is attained, wherein *I*_Ca_ and *I*_K_ counterbalance, while deactivation of Na_v_1.5 ensures *I*_Na_ is negligible (step 4 - Figs. 8a,c). While *c*_i_ decays from its peak, *I*_Ca_ continues to increase—owing to the large value of 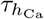—which eventually results in *I*_Ca_ dominating *I*_K_ and concomitant depolarisation (step 5 - Figs. 8a-c). Continued depolarisation leads to the inactivation of Ca_v_3.2, activation of Na_v_1.5, and an increase in *I*_K_ (step 6 - Fig. 8c). The cell then returns to a state of depolarised equilibrium (step 1). A complementary way of visualising these steps is afforded by the trajectory traced out by the system in the *c*_i_-*V*_m_ plane (Fig. 8d).

HVTs are observed for values of *σ* in the range 0.02 ms^*−*1*/*2^ ≲ *σ* ≲ 0.2 ms^*−*1*/*2^. For values of *σ* within this range that are smaller than 0.05 ms^*−*1*/*2^ (the value used in Fig. 6), longer lasting HVTs can occur—see the HVT between 20 - 40 s in Fig. 9a which occurs for *σ* = 0.04 ms^*−*1*/*2^. Interestingly, HVTs of variable durations were observed via voltage imaging (Quicke et al. 2022). Note that the peak value of *c*_i_ during the HVT between 20 - 40 s (Fig. 9b) of ∼ 900 nM is considerably greater than the hyperpolarised steady state value of 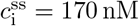 (see Section 3.1). Peak values of *c*_i_ that are comparable—or even an order of magnitude larger—are sometimes seen accompanying simulated HVTs. While such high values are physiologically unrealistic—an uninhibited ‘runaway process’ occurring due to the multiplicative noise term used to model CICR—their occurrence does not alter the qualitative nature of the accompanying HVTs, compared to those in Fig. 6a, for instance.

**Figure 9:**
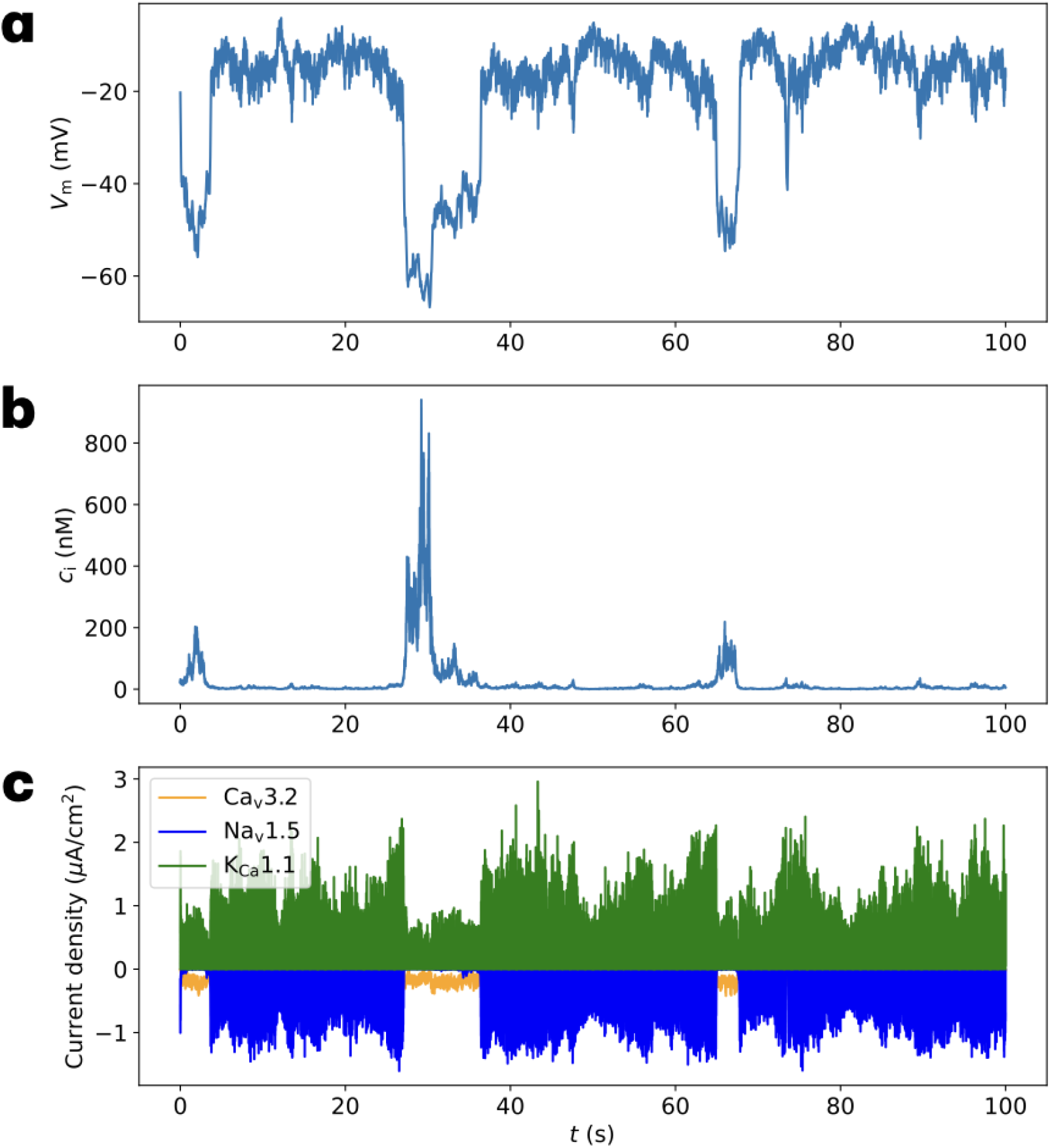
A 100 s simulation of the SDEs for reduced *σ* displaying HVTs. Stochastic version of Eq. 14 and Eq. 18 simulated for *σ* = 0.04 ms^*−*1*/*2^, with corresponding time series for *V*_m_ **(a)**, *c*_i_ **(b)** and currents for each ionic species **(c)**. Note that the duration of the HVT centred at ∼ 30 s is roughly twice that of the longest lasting HVTs in Fig. 6, which were generated for *σ* = 0.05 ms^*−*1*/*2^.

As *σ* is increased toward 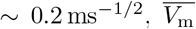 increases while 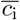 decreases; past ∼ 0.2 ms^*−*1*/*2^, HVTs are no longer observed (Fig. 10). We see that transient hyperpolarising fluctuations occur (Fig. 10a) with concomitant spikes in *c*_i_ (Fig. 10b). However, such fluctuations are distinct from HVTs discussed above: an examination of the ionic currents (Fig. 10c) reveals that noisy variations in *I*_K_ and *I*_Na_ are entirely responsible for these events, while *I*_Ca_ is negligible throughout the simulation. The unreasonably high and low values of 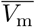 and 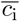, respectively—seen in Fig. 10 for instance—suggests that values of *σ*, above the range identified for HVTs, cannot capture physiologically realistic values of *V*_m_ and *c*_i_.

**Figure 10:**
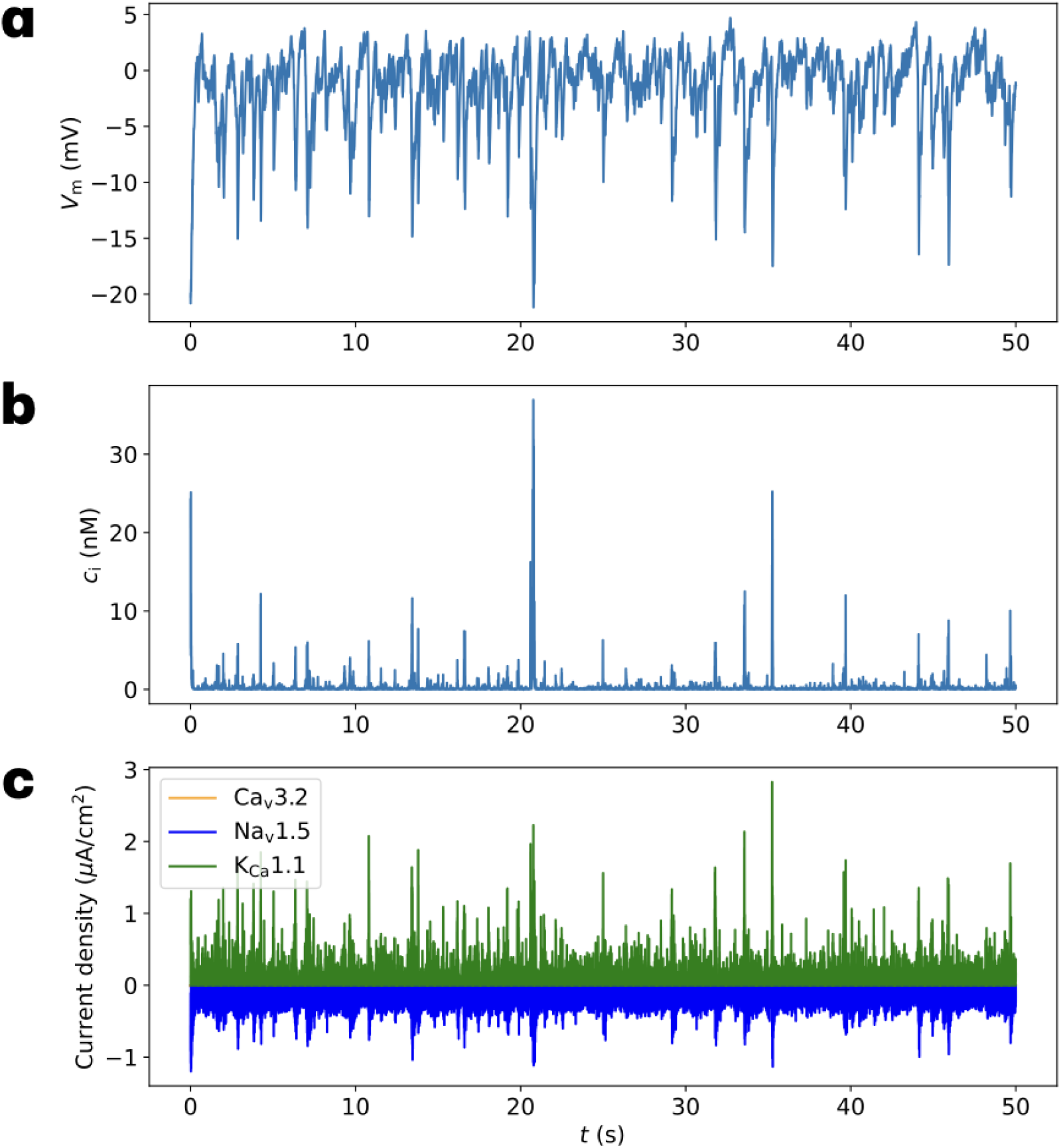
A 50 s simulation of the SDEs for increased *σ*. Stochastic version of Eq. 14 and Eq. 18 simulated for *σ* = 0.3 ms^*−*1*/*2^, with corresponding time series for *V*_m_ **(a)**, *c*_i_ **(b)** and currents for each ionic species **(c)**. The average values of *V*_m_ and *c*_i_ are 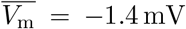 mV and 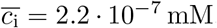 mM, respectively. The transient hyperpolarising fluctuations in **(a)** have a different character from the HVTs seen in, say, Fig. 6a, which is reflected in the contrasting current time series (compare **(c)** with Fig. 6c).

### 3.3 Inhibition of CICR leads to hyperpolarisation

As discussed in Section 3.2, there is a lower limit for *σ* below which HVTs do not occur. Reducing *σ* below this limit simulates the effect of inhibiting CICR, and it is thus necessary to understand the system dynamics under this perturbation. Let us consider the value of *σ* = 2 · 10^*−*3^ ms^*−*1*/*2^, which is a factor of 25 smaller than the value employed for the simulation in Fig. 6. The *V*_m_ time series (Fig. 11a) is characteristic of the dynamics seen for such reduced values of *σ*: stochastic fluctuations of the ion channels hyperpolarise *V*_m_, at first gradually, then abruptly, on the timescale of ∼ 1 s. This is followed by a slower relaxation to ∼ −55 mV. After an initial transient of ∼ 1 s, the *c*_i_ time series (Fig. 11b) displays noisy exponential relaxation toward the equilibrium value of 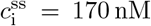 for the hyperpolarised steady state (see Section 3.1). This relaxation occurs with a characteristic timescale of *γ*^*−*1^ = 10 s, as can be seen from Eq. 17 if *I*_Ca_ is approximately constant (see *I*_Ca_ time series in Fig. 11c after ∼ 1 s). Note the contrast between the timescale for *c*_i_ equilibration when CICR is inhibited, and the timescale for *c*_i_ to reach its peak value (and subsequently decay) over the course of a spike (of the order 100 ms), when CICR is active—see Fig. 8b. Similar behaviour to that described above for *σ* = 2 · 10^*−*3^ ms^*−*1*/*2^ is seen for higher values of *σ*, except that the relaxation to equilibrium of *V*_m_ and *c*_i_ is noisier.

**Figure 11:**
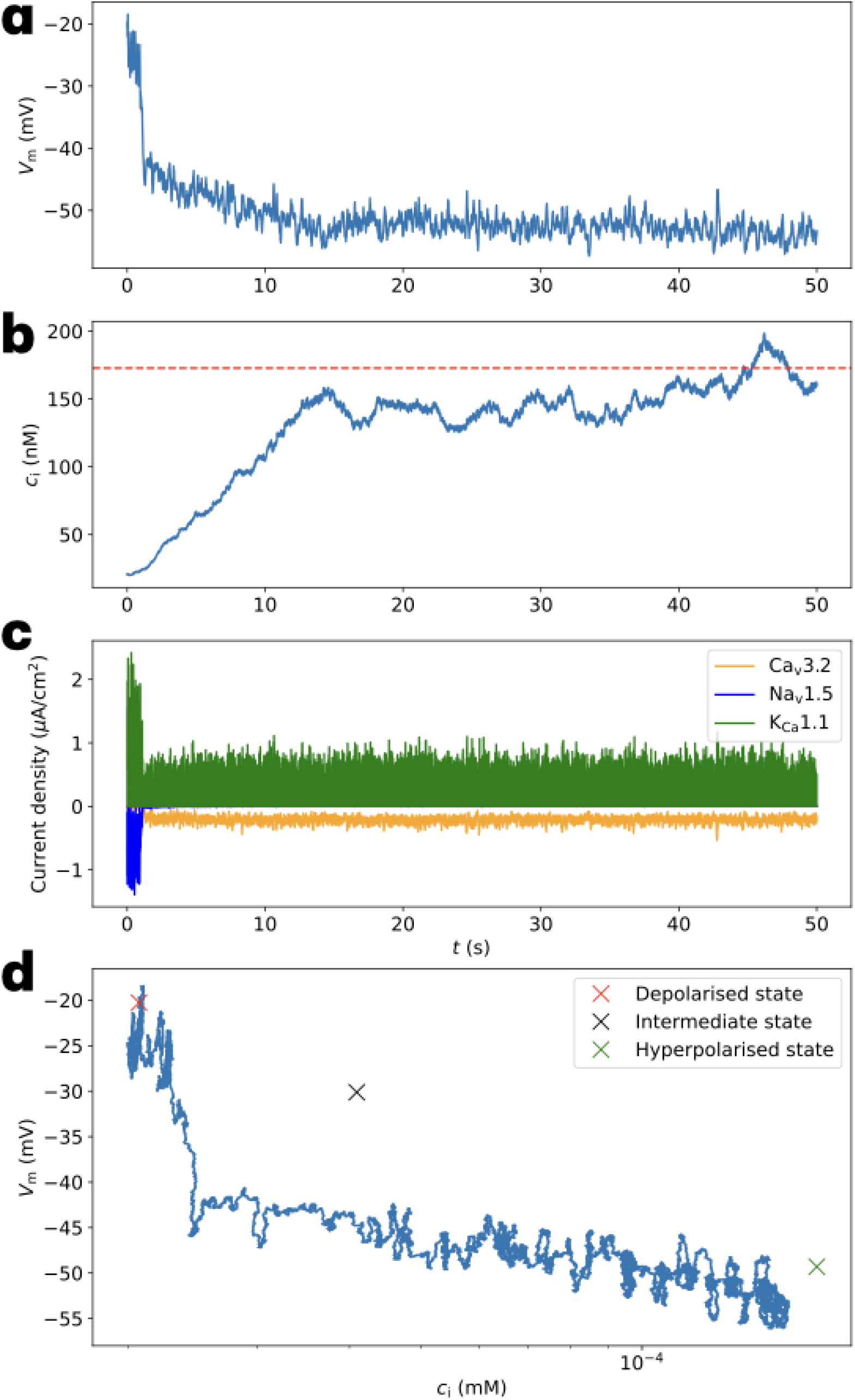
A simulation of the SDEs for much reduced *σ*, displaying slow relaxation to hyperpolarised equilibrium. Stochastic version of Eq. 14 and Eq. 18 simulated for 50 s for *σ* = 2 · 10^*−*3^ ms^*−*1*/*2^, with corresponding time series for *V*_m_ **(a)**, *c*_i_ **(b)** and currents for each ionic species **(c)**. The dashed horizontal line in **(b)** marks 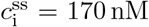 nM for the hyperpolarised steady state. **(d)** Trajectory of the system in the *c*_i_-*V*_m_ plane during the first 15 s of the simulation.

It is illuminating to consider the trajectory of the system in the *c*_i_-*V*_m_ plane under inhibition of CICR (Fig. 11d). The slow relaxation of the system toward the vicinity of the hyperpolarised equilibrium (after the abrupt hyperpolarisation) can be seen clearly. Note that in both Fig. 11d and Fig. 8d—for which CICR is active—the system reaches similar regions of the *c*_i_-*V*_m_ state space, (∼ 5 · 10^*−*5^ mM, ∼ − 45 mV). Despite this, the subsequent evolution of the system for the two cases is markedly different. When CICR is inhibited, the system is ‘pulled toward’ the hyperpolarised state, which functions as an attracting state. On the other hand, when CICR is active, the system appears to be ‘repelled’ by the hyperpolarised state, as it returns to depolarised equilibrium. Under inhibition of CICR, the system reaches (∼ 5 · 10^*−*5^ mM, ∼ −45 mV) in ∼ 5 s, which is ample time for all the gating variables to equilibrate (note from Fig. 11c that at ∼ 5 s, *I*_K_ and *I*_Ca_ have effectively equilibrated). As a result, the vectors of the phase plane, which ‘pave’ trajectories toward hyperpolarised equilibrium—analogous to Fig. 5a—guide the system to this end. In contrast, during a HVT (for which CICR activation is required), the time between the start (step 1) and the system reaching (∼ 5 · 10^*−*5^ mM, ∼ −45 mV) (step 3) is ∼ 200 ms (Figs. 8a,b). This is not enough time for *h*_Ca_ to equilibrate—note from Fig. 8c how |*I*_Ca_| is still rising at step 3. In conjunction with the stochastic dynamics of *c*_i_ (due to multiplicative noise), this results in the system not relaxing toward the hyperpolarised state ^19^.

### 3.4 Blockage of ion channels

We can probe the effect of pharmacological inhibition of ion channels in our model by reducing the maximum channel conductances, and thus, the number of functionally active ion channels. As mentioned in Section 2.1.4, the SDEs derived by Fox and Lu—which we have employed thus far—approximate Markov processes to model sparse ion channel expression. Studies have revealed that the SDE approximation can yield inaccuracies relative to ‘exact’ Markov process models—for instance in AP firing statistics—that persist even for increasing numbers of ion channels (Bruce 2009; Goldwyn and Shea-Brown 2011). Given this, and the low numbers of ion channels that we have estimated, a further reduction in these numbers (to simulate ion channel blockage) no longer permits us to integrate our model SDEs to explore system dynamics. Despite this, we can still gain useful information about the effect of ion channel blockage by examining the stability landscape of the system.

Let us consider the effect of reducing the maximum conductances in turn by a factor of 10, relative to their values in Table 1, while keeping all other parameters unchanged. Figs. 12a,b display the effect of reducing 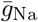 on the system’s stability. We see a dramatic change from Fig. 4, where bistability gives way to a single hyperpolarised steady state (Fig. 12a). An examination of the system’s phase plane in the vicinity of this state (Fig. 12b) reveals that it is a steady state, and thus the system is now monostable. Similarly, reduction of 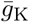 leads to monostability: a single depolarised steady state (Figs. 12c,d). The emergence of monostability after sufficiently reducing 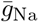 or 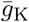 is consistent with observations that application of TTX or IbTX—which specifically block Na_v_1.5 or K_Ca_1.1, respectively—suppresses *V*_m_ fluctuations and HVTs in MDA-MB-231 cells (Quicke et al. 2022).

**Figure 12:**
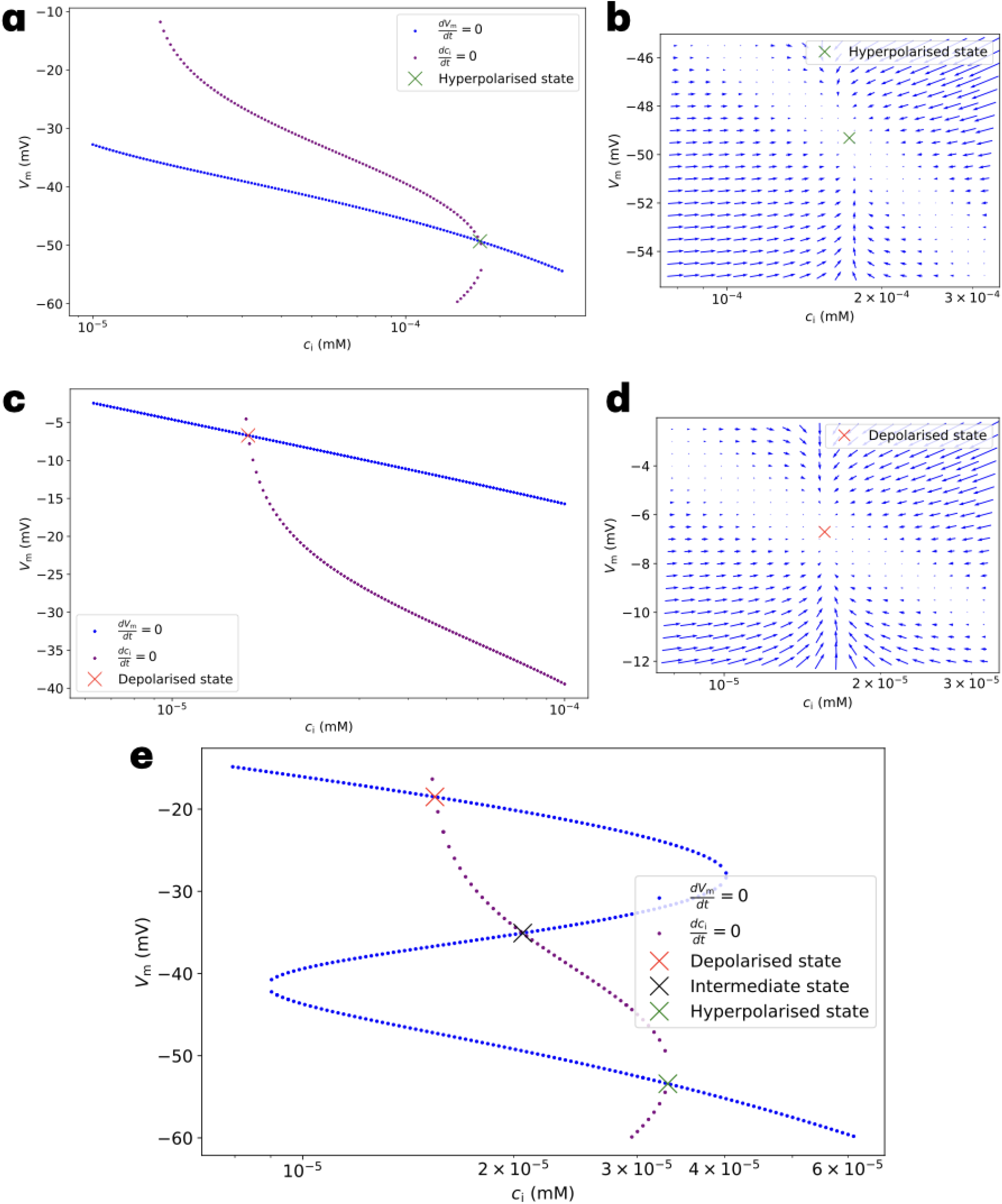
Stability landscapes of the system after reduction of the maximum conductance densities. The effect of 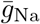 (top row), 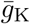 (middle row) and 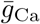 (bottom row) being reduced by a factor of 10. **(a)**,**(c)**,**(e)** Nullclines for *c*_i_ (purple dots) and *V*_m_ (blue dots). The points of intersection of the nullclines (marked by crosses) correspond to the steady states of the system. In **(e)**, bistability persists since the nature of stability of the three equilibria are unchanged. **(b)**,**(d)** Examination of the system’s phase planes in the neighbourhood of the steady states confirms that they are stable. For clarity, 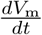 is scaled by 10^*−*3^, which does not alter the nature of stability of the hyperpolarised or depolarised state.

Interestingly, while reduction of 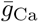 does not remove bistability, quantitative changes in the steady states occur, which are especially pronounced for the hyperpolarised state (Fig. 12e) ^20^. For this state, 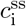 decreases appreciably from its prior value of 1.7·10^*−*4^ mM to 3.3·10^*−*5^ mM, along with a small hyperpolarisation of 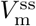. The inability to simulate the system dynamics now makes definitive prediction of the effect of Ca_v_3.2 blockage not possible. Nonetheless, two plausible outcomes are: (i) a slow relaxation to hyperpolarised equilibrium (similar to inhibition of CICR) due to the reduction in the range of *c*_i_ in the stability landscape, and the correspondingly diminished effect of the multiplicative noise term which captures CICR; (ii) the continued occurrence of HVTs, albeit with *c*_i_ spikes of much reduced amplitude.

## 4 Discussion

The main conclusions of this study are as follows:

1. A minimal HH–type model—based on Na_v_1.5, Ca_v_3.2, and K_Ca_1.1 ion channels coupled to Ca^2+^ influx and clearance—is sufficient to predict the emergence of *V*_m_ bistability in MDA-MB-231 cells.
2. The inclusion of stochasticity in the dynamics of *c*_i_ and *V*_m_—reflecting CICR and low density ion channel expression, respectively—induces transitions between the depolarised and hyper-polarised steady states. Such HVTs are in agreement with experimental observations, with the predicted |Δ*V*_m_| ≃ 30 mV being comparable to the changes recorded experimentally. Our model predicts that specific blockage of Na_v_1.5 or K_Ca_1.1 would lead to monostability, and thus the disappearance of HVTs, again, consistent with voltage imaging.
3. The model contains a single adjustable parameter, *σ*, which reflects the strength of CICR. Our simulations yield HVTs for a range of values of *σ*, which controls some of the characteristics of HVTs, for instance, their duration.
4. A novel prediction of our model is that inhibition of CICR—simulated by a sufficient reduction of *σ*—would induce hyperpolarisation, and thus suppress HVTs.

### 4.1 Hyperpolarisation driven Ca^2+^ transient signalling via T-type VGCCs

Voltage imaging has uncovered a new window through which to measure, for the first time, the rich and varied ‘dynamic electrical signatures’ of individual breast cancer cells, including those of the MDA-MB-231 cell line (Quicke et al. 2022). The modelling of such dynamics—as we have presented in this work—thus sheds new light on long-standing questions regarding signalling mechanisms in MDA-MB-231 cells, and may have implications for electrically active cells more generally. For instance, as noted in (Gray et al. 2004), it has been widely accepted that Ca^2+^ entry in non-excitable cancer cells cannot be regulated by VGCCs, unless VGCC opening occurs via depolarisation over the course of the cell cycle (Panner and Wurster 2006). In contrast, our model proposes that during a HVT, recovery from inactivation of Ca_v_3.2 channels enables Ca^2+^ influx, and a subsequent cytosolic Ca^2+^ transient, fuelled by CICR. This influx of Ca^2+^ is an example of the ‘window current’ characteristic of T-type VGCCs, which was shown previously to induce *V*_m_ bistability within thalamic neurons (see (Crunelli et al. 2005) and references therein). Interestingly, steady state window currents through Ca_v_3.2 channels have been hypothesised previously for breast cancer (Taylor et al. 2008) and prostate cancer cells (Gackiere et al. 2013; Prevarskaya, Skryma, and Shuba 2018). However, it appears that the proposed ability of VGCCs to transduce a *V*_m_ hyperpolarisation signal into a cytosolic Ca^2+^ transient is potentially novel, and complements their well known ability to transduce AP signals to Ca^2+^ signals (Catterall 2011). This suggests that K_Ca_1.1-Ca_v_3.2 channel complexes may form in MDA-MB-231 cells (as in prostate cancer cells (Gackiere et al. 2013)), and could play a significant role in regulating tumourigenic processes, such as proliferation (Guéguinou et al. 2014).

### 4.2 Channel noise due to sparse ion channel expression

While aberrant ion channel expression has been detected in cancer cells for some time now (Prevarskaya, Skryma, and Shuba 2018), the functional consequences of low density channel expression in the cell membrane have, to the best of our knowledge, been unexplored thus far. Here, our estimates of this density (from prior studies) for Na_v_1.5, Ca_v_3.2 and K_Ca_1.1 channels in MDA-MB-231 cells, suggest that these channels are sparsely expressed compared to ‘key’ channels in typical neurons, which necessitates modelling *V*_m_ stochastically. Importantly, stochastic fluctuations which decrease the Na_v_1.5 channel conductance are seen to be the initiating step of a HVT (Fig. 7); taking the deterministic limit (by assuming ‘large’ numbers of ion channels, *N*_X_→ ∞ in Eq. 16) would lead to the disappearance of this mechanism. Further implications of channel noise in isolated and interacting cancer cells await to be uncovered, and, with noise being an inescapable feature of the nervous system (Faisal, Selen, and Wolpert 2008), cancer neuroscience appears to be an intriguing arena for future studies of this phenomenon (Mancusi and Monje 2023; Winkler et al. 2023).

### 4.3 Stochastic modelling of Ca^2+^ spiking

In this study, we modelled the positive feedback nature of CICR using multiplicative noise, which generates cytosolic Ca^2+^ spikes. Such spikes are understood to vary smoothly in time—and are thus modelled deterministically—with spiking emerging from the interaction of stochastic processes at the level of IP_3_Rs or RyRs (Dupont, Combettes, et al. 2011; Dupont, Falcke, et al. 2016). Although our phenomenological modelling of Ca^2+^ spiking is thus a caricature of this process—being inherently noisy—it nonetheless captures the essential features of a fast positive feedback process followed by a slower negative feedback process. Such a caricature comes at the cost of simulations sometimes yielding unreasonably high peak values of *c*_i_, as we noted in Section 3.2 ^21^. However, we think this is acceptable, given that our motivation here is to explain, in a minimal fashion, *V*_m_ bistability and HVTs observed in MDA-MB-231 cells. The unrealistic values of *c*_i_ do not alter the realistic mechanisms underlying bistability and HVTs that emerge from our model. Furthermore, despite its simplicity, the stochastic modelling of Ca^2+^ dynamics yields the testable prediction that inhibition of CICR—via silencing or inhibition of IP_3_R3 (RyR) (using e.g. Xestospongin C (Ryanodine))—results in hyperpolarisation. Simultaneous Ca^2+^ and *V*_m_ imaging experiments could confirm the occurrence of Ca^2+^ spikes and their predicted synchrony with HVTs.

## 5 Limitations of the study

We chose the ion channels utilised in our model given the robust evidence for their expression in MDA-MB-231 cells. However, the assumed dynamics of these ion channels (based on measured time constants etc.) stems from studies on non-cancerous cells, for instance, K_Ca_1.1 expressed in bullfrog sympathetic ganglion cells (Yamada, Koch, and Adams 1989). Indeed, it is well known that alternative splicing enables different tissues to express structurally and functionally distinct isoforms of the same ion channel. It is almost inevitable therefore, that the dynamics of the ion channels expressed in MDA-MB-231 cells could differ from what we have assumed in our model. Despite this, and as long as these differences are not substantial, they would only affect the predictions of our model quantitatively, and leave the qualitative aspects unchanged.

In Section 3.4, we highlighted the limitations of the SDE approximation of Markov processes, which prevents us from integrating the model SDEs to explore dynamics under the effect of ion channel blockage. Indeed, inaccuracies in the prediction of AP firing statistics (compared to Markov chain ion channel models) also imply that a quantitative comparison of the frequency of simulated and experimentally observed HVTs would not be viable. Interestingly, Fox and Lu derived an alternative system of high-dimensional SDEs which enables one to overcome such issues, albeit at a greater computational cost than the SDE approximation employed here (Fox and Lu 1994; Goldwyn and Shea-Brown 2011). This alternative system models the dynamics of the fraction of ion channels in distinct configurations, and its predictions are on a par with those of Markov process models (see (Goldwyn and Shea-Brown 2011) and references therein). Thus, the use of higher-dimensional SDEs could form the basis of future modelling extending the work presented here.

## 6 Future perspectives

Since its introduction, the HH formalism has been used to model electrical activity in diverse cell types beyond neurons, including cardiac pacemaker cells (Noble 1960), pancreatic *β*-cells (Chay and Keizer 1983), and pituitary somatotrophs (Tsaneva-Atanasova et al. 2007). Meanwhile, over the past decade or so, the role of *V*_m_ bistability in biophysical models of tumour dynamics—as well as in the contexts of development and regeneration—has been investigated (Cervera, Levin, and Mafe 2023) ^22^. Motivated by voltage imaging observations (Quicke et al. 2022), this study proposes that the coupling of the stochastic HH formalism to cytosolic Ca^2+^ dynamics yields bistability within the *c*_i_-*V*_m_ state space, and noise induced switching between the stable states. It is increasingly well understood that tumourigenic processes are dependent on pathways regulated by Ca^2+^ signalling, which is often dysregulated or remodelled (Monteith, Prevarskaya, and Roberts-Thomson 2017). The relationship between HVTs and Ca^2+^ spiking and remodelled Ca^2+^ signalling remains to be uncovered in the future.

The MDA-MB-231 cell line—which has been used extensively in cancer research—has been shown to display both genetic and phenotypic heterogeneity (Martín-Pardillos et al. 2019; Hapach et al. 2021). Such heterogeneity may partially explain why only ∼ 7% of the cells examined in (Quicke et al. 2022) displayed *V*_m_ fluctuations, and similarly, why prior studies observed only a subpopulation of cells expressing functional VGSCs (Fraser et al. 2005). Furthermore, Quicke et al. uncovered distinct dynamic electrical signatures (classes of fluctuations) across the entire population of cells observed, including ‘blinking’ and ‘waving’ patterns. The HVTs generated by our model would appear to fall into the (large) blinking class; how different classes of fluctuations arise—and the possible role that cell line heterogeneity may play in this—is a topic for future study. Similarly, the methodology outlined here could potentially be employed, with modifications, to explain the electrical activity of other cancer cell lines. In particular, analysing cancer cells with varying metastatic ability (known to involve different ion channels) would be extremely worthwhile.

This work has focused on the *V*_m_ dynamics of individual MDA-MB-231 cells, and a natural next step would be to investigate such dynamics for cellular collectives. Indeed, evidence of intercellular signalling has been uncovered, by way of temporal correlations between different cells imaged simultaneously, for instance, a propagating wave of HVTs (Quicke et al. 2022). In fact, prior to the work of Quicke et al., the possibility of intercellular Ca^2+^ waves mediating signalling within a cohort of MDA-MB-231 cells was proposed (Ribeiro et al. 2020). Propagating waves of HVTs would seem to imply a deterministic ‘triggering’ of a HVT, from one cell to another, and contrasts with the mechanism of gating fluctuations yielding stochastic HVTs, proposed here. Robust verification of intercellular Ca^2+^ signalling would suggest the need to incorporate other elements of the Ca^2+^ signalling ‘toolbox’ (Dupont, Falcke, et al. 2016) to amend or extend our model.

In summary, we have shown that a minimal, stochastic HH–type model—consisting of Na_v_1.5, Ca_v_3.2, and K_Ca_1.1 ion channels coupled to intracellular Ca^2+^ dynamics incorporating CICR—naturally gives rise to *V*_m_ bistability in MDA-MB-231 cells. Noise-driven transitions between the depolarised and hyperpolarised steady states qualitatively capture HVTs observed experimentally. The stability landscape of the system implies that pharmacological blockage of Na_v_1.5 or K_Ca_1.1 should abolish bistability and HVTs, in agreement with voltage imaging observations. Furthermore, simulations yield the novel prediction that inhibition of CICR would shift cells into a persistently hyperpolarised state, preventing HVTs. Beyond capturing key experimental observations for the MDA-MB-231 cell line, our model provides a novel framework in which sparse ion channel expression and cytosolic Ca^2+^ dynamics can regulate the dynamic electrical behaviour of breast cancer cells. Thus, this approach could help our quantitative understanding of the pathophysiological role of bioelectrical signalling in cancer cells. Importantly, manipulating *V*_m_ has been shown to be an effective means of controlling cancer (B. Chernet and Levin 2013; B. T. Chernet et al. 2016). Accordingly, determining the precise role of ion channels in the behavior of *V*_m_ could ultimately enable novel therapies involving repurposed ion channel drugs (Kofman and Levin 2024).

## Abbreviations

AP: Action potential
Ca_v_3.2: Voltage-gated calcium channel subtype 3.2
CICR: Calcium-induced calcium release
ER: Endoplasmic reticulum
HVT: Hyperpolarising voltage transient
HH: Hodgkin-Huxley
IbTX: Iberiotoxin
IP_3_R: Inositol (1,4,5)-trisphosphate receptor
IP_3_R3: Type 3 inositol (1,4,5)-trisphosphate receptor
K_Ca_1.1: Voltage- and calcium-activated potassium channel subtype 1.1
Na_v_1.5: Voltage-gated sodium channel subtype 1.5
ODE: Ordinary differential equation
RyR: Ryanodine receptor
SDE: Stochastic differential equation
TTX: Tetrodotoxin
VGCC: Voltage-gated calcium channel
VGSC: Voltage-gated sodium channel
V_m_: Membrane potential

## Acknowledgements

We are grateful to Dr Amanda Foust for commenting on the manuscript.

## CRediT authorship contribution statement

**Suvabrata De:** Conceptualisation, Methodology, Software, Formal analysis, Writing – original draft. **Mustafa B. A. Djamgoz:** Conceptualisation, Writing – review & editing.

## Data availability

Data sharing is inapplicable to this study as no datasets were generated or analysed.

## Code availability

The Python code employed in this study is available from SD upon reasonable request.

## Footnotes

1 From here on, we refer to the neonatal form of Na_v_1.5 as simply Na_v_1.5.

2 Time is expressed in units of ms throughout.

3 We assume that the deactivation time constant data presented in Fig. 1F of (Zhang et al. 2013) provides a useful approximation for the activation time constant employed in our HH formalism. Additionally, we assume that the peak inactivation time constant in Fig. 1F of (Zhang et al. 2013) is reached at -50 mV, and thus ‘extrapolate symmetrically’ for time constants below -50 mV.

4 We express current densities in units of *µ*A cm^*−*2^ throughout, and for brevity take ‘current’ to be synonymous with ‘current density’ from here on.

5 A variable Nernst potential is necessary given that *c*_i_ varies by orders of magnitude in our model: *E*_Ca_ ranges between 96-154 mV for *c*_i_ ranging between 10^*−*3^-10^*−*5^ mM.

6 We use the room temperature value of 295.15 K (22°C), in accordance with the values and methods quoted by the studies we use to model ion channel dynamics.

7 We estimate the ionised concentration of calcium to be half of the total concentration of 10 mM quoted in (Taylor et al. 2008).

8 Blockage of K_Ca_1.1 in both studies diminished the corresponding phenomena.

9 The rate constants for opening (forward) and closing (backward) of the channel.

10 From Eq. 11a, for *V*_m_ = 50 mV and assuming *c*_i_ = 10^*−*4^ mM, it is found that 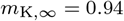.

11 (Kang et al. 2021; Selimi et al. 2023)

12 (Engbers, Zamponi, and Turner 2013)

13 (Contreras et al. 2013)

14 A value of 2 10^*−*5^ mM (quoted as ‘*c*_0_’) is utilised for *c*_rest_ in prior studies modelling intracellular Ca^2+^ dynamics (Rüdiger, Shuai, and Sokolov 2010; Dobramysl, Sten Rüdiger, and Erban 2016).

15 The first term on the right-hand side of Eq. 17 converts current flux into concentration flux, since 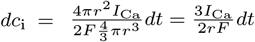, which has units of nmol cm^*−*3^=*µ*M (the factor of 2 reflects the valence of Ca^2+^ ions and recall that *F* ≡ *eN*_A_). The additional factor of 1000 in Eq. 17 is needed to convert to mM units, which is the unit of concentration we employ.

16 For instance, in the original HH model, at the peak of an AP 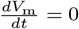 instantaneously, but not indefinitely.

17 These curves are known as the *V*_m_- and *c*_i_-nullclines, respectively.

18 Strictly speaking, the phase planes in Fig. 5 faithfully represent the system dynamics if the gating variables are at, or close to, their equilibrium (steady state) values. In Section 3.3, we discuss that if the gating variables have not equilibrated, the system dynamics can be very different from that suggested by the phase plane.

19 We note that when the system is initialised at (5 10^*−*5^ mM, 45 mV) with the gating variables equilibrated and CICR is active (*σ* = 0.05 ms^*−*1*/*2^), relaxation to hyperpolarised equilibrium does not occur. In other words, equilibration of the gating variables does not guarantee relaxation.

20 The phase planes of Fig. 5 are still a qualitatively valid description of the stability of the equilibrium points now.

21 The runaway process could be dampened by replacing the stochastic term in Eq. 18 with (for example) 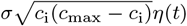, at the expense of introducing an extra model parameter, *c*_max_.

22 See references in (Cervera, Levin, and Mafe 2023) for numerous examples of *V*_m_ bistability in different cell types.

## References

Abdul, Mansoor, Sian Ramlal, and Naseema Hoosein (2008). “Ryanodine receptor expression correlates with tumor grade in breast cancer”. In: Pathology & Oncology Research 14, pp. 157– 160.

Awad, Mohamad Fawzi et al. (2024). “Dielectrophoretic and electrochemical impedance mapping of metastatic potential in MDA-MB-231 breast cancer cells using inkjet-printed castellated microarray”. In: Lab on a Chip 24.18, pp. 4264–4274.

Bhargava, Anamika and Sumit Saha (2019). “T-Type voltage gated calcium channels: a target in breast cancer?” In: Breast cancer research and treatment 173, pp. 11–21.

Bruce, Ian C (2009). “Evaluation of stochastic differential equation approximation of ion channel gating models”. In: Annals of biomedical engineering 37, pp. 824–838.

Carvalho, Joao (2021). “A bioelectric model of carcinogenesis, including propagation of cell membrane depolarization and reversal therapies”. In: Scientific Reports 11.1, p. 13607.

Catterall, William A (2011). “Voltage-gated calcium channels”. In: Cold Spring Harbor perspectives in biology 3.8, a003947.

Cervera, Javier, Antonio Alcaraz, and Salvador Mafe (2016). “Bioelectrical signals and ion channels in the modeling of multicellular patterns and cancer biophysics”. In: Scientific reports 6.1, p. 20403.

Cervera, Javier, Michael Levin, and Salvador Mafe (2023). “Bioelectricity of non-excitable cells and multicellular pattern memories: Biophysical modeling”. In: Physics reports 1004, pp. 1–31.

Chay, Teresa Ree and Joel Keizer (1983). “Minimal model for membrane oscillations in the pancreatic beta-cell”. In: Biophysical journal 42.2, pp. 181–189.

Chernet, Brook and Michael Levin (2013). “Endogenous voltage potentials and the microenvironment: bioelectric signals that reveal, induce and normalize cancer”. In: Journal of clinical & experimental oncology, S1–002.

Chernet, Brook T et al. (2016). “Use of genetically encoded, light-gated ion translocators to control tumorigenesis”. In: Oncotarget 7.15, p. 19575.

Contreras, Gustavo F et al. (2013). “A BK (Slo1) channel journey from molecule to physiology”. In: Channels 7.6, pp. 442–458.

Crunelli, Vincenzo et al. (2005). “The ‘window’T-type calcium current in brain dynamics of different behavioural states”. In: The Journal of physiology 562.1, pp. 121–129.

Di Francesco, Dario and Denis Noble (1985). “A model of cardiac electrical activity incorporating ionic pumps and concentration changes”. In: Philosophical Transactions of the Royal Society of London. B, Biological Sciences 307.1133, pp. 353–398.

Djamgoz, Mustafa BA (2024). “Electrical excitability of cancer cells—CELEX model updated”. In: Cancer and Metastasis Reviews, pp. 1–13.

Dobramysl, Ulrich, Sten Rüdiger, and Radek Erban (2016). “Particle-based multiscale modeling of calcium puff dynamics”. In: Multiscale Modeling & Simulation 14.3, pp. 997–1016.

Dupont, Geneviéve, Laurent Combettes, et al. (2011). “Calcium oscillations”. In: Cold Spring Harbor perspectives in biology 3.3, a004226.

Dupont, Geneviéve, Martin Falcke, et al. (2016). Models of calcium signalling. Vol. 43. Springer.

Endo, Makoto (2009). “Calcium-induced calcium release in skeletal muscle”. In: Physiological reviews 89.4, pp. 1153–1176.

Engbers, Jordan DT, Gerald W Zamponi, and Ray W Turner (2013). “Modeling interactions between voltage-gated Ca2+ channels and KCa1. 1 channels”. In: Channels 7.6, pp. 524–529.

Fabiato, Alexandre (1983). “Calcium-induced release of calcium from the cardiac sarcoplasmic reticulum”. In: American Journal of Physiology-Cell Physiology 245.1, pp. C1–C14.

Faisal, A Aldo, Luc PJ Selen, and Daniel M Wolpert (2008). “Noise in the nervous system”. In: Nature reviews neuroscience 9.4, pp. 292–303.

Fox, Ronald F and Yan-nan Lu (1994). “Emergent collective behavior in large numbers of globally coupled independently stochastic ion channels”. In: Physical Review E 49.4, p. 3421.

Fraser, Scott P et al. (2005). “Voltage-gated sodium channel expression and potentiation of human breast cancer metastasis”. In: Clinical cancer research 11.15, pp. 5381–5389.

Gackiere, Florian et al. (2013). “Functional coupling between large-conductance potassium channels and Cav3. 2 voltage-dependent calcium channels participates in prostate cancer cell growth”. In: Biology open 2.9, pp. 941–951.

Goldwyn, Joshua H and Eric Shea-Brown (2011). “The what and where of adding channel noise to the Hodgkin-Huxley equations”. In: PLoS computational biology 7.11, e1002247.

Gray, Lloyd S et al. (2004). “The role of voltage gated T-type Ca2+ channel isoforms in mediating “capacitative” Ca2+ entry in cancer cells”. In: Cell Calcium 36.6, pp. 489–497.

Groff, Jeffrey R, Hilary DeRemigio, and Gregory D Smith (2009). “Markov chain models of ion channels and calcium release sites”. In: Stochastic Methods in Neuroscience. Ed. by Gabriel J Lord and Carlo Laing. Oxford Univ. Press. Chap. 2, pp. 29–64.

Guéguinou, Maxime et al. (2014). “KCa and Ca2+ channels: the complex thought”. In: Biochimica et Biophysica Acta (BBA)-Molecular Cell Research 1843.10, pp. 2322–2333.

Han, Arum, Lily Yang, and A Bruno Frazier (2007). “Quantification of the heterogeneity in breast cancer cell lines using whole-cell impedance spectroscopy”. In: Clinical cancer research 13.1, pp. 139–143.

Hapach, Lauren A et al. (2021). “Phenotypic heterogeneity and metastasis of breast cancer cells”. In: Cancer research 81.13, pp. 3649–3663.

Higham, Desmond and Peter Kloeden (2021). An introduction to the numerical simulation of stochastic differential equations. SIAM.

Hodgkin, Alan L and Andrew F Huxley (1952). “A quantitative description of membrane current and its application to conduction and excitation in nerve”. In: The Journal of physiology 117.4, p. 500.

Kang, Po Wei et al. (2021). “Elementary mechanisms of calmodulin regulation of NaV1. 5 producing divergent arrhythmogenic phenotypes”. In: Proceedings of the National Academy of Sciences 118.21, e2025085118.

Keener, James and James Sneyd (2008). Mathematical Physiology: I: Cellular Physiology. Springer Science & Business Media.

Khaitan, Divya et al. (2009). “Role of KCNMA1 gene in breast cancer invasion and metastasis to brain”. In: BMC cancer 9, pp. 1–11.

Kofman, Karina and Michael Levin (2024). “Bioelectric pharmacology of cancer: a systematic review of ion channel drugs affecting the cancer phenotype”. In: Progress in biophysics and molecular biology.

Levin, Michael (2021). “Bioelectric signaling: Reprogrammable circuits underlying embryogenesis, regeneration, and cancer”. In: Cell 184.8, pp. 1971–1989.

Li, Yue-Xian and John Rinzel (1994). “Equations for InsP3 receptor-mediated [Ca2+] i oscillations derived from a detailed kinetic model: a Hodgkin-Huxley like formalism”. In: Journal of theoretical Biology 166.4, pp. 461–473.

Mancusi, Rebecca and Michelle Monje (2023). “The neuroscience of cancer”. In: Nature 618.7965, pp. 467–479.

Martín-Pardillos, Ana et al. (2019). “The role of clonal communication and heterogeneity in breast cancer”. In: BMC cancer 19, pp. 1–26.

Monteith, Gregory R, Natalia Prevarskaya, and Sarah J Roberts-Thomson (2017). “The calcium– cancer signalling nexus”. In: Nature Reviews Cancer 17.6, pp. 373–380.

Morley Sinéad T, Michael T Walsh, and David T Newport (2017). “The advection of microparticles, MCF-7 and MDA-MB-231 breast cancer cells in response to very low Reynolds numbers”. In: Biomicrofluidics 11.3.

Mound, Abdallah, Lise Rodat-Despoix, et al. (2013). “Molecular interaction and functional coupling between type 3 inositol 1, 4, 5-trisphosphate receptor and BKCa channel stimulate breast cancer cell proliferation”. In: European journal of cancer 49.17, pp. 3738–3751.

Mound, Abdallah, Alexia Vautrin-Glabik, et al. (2017). “Downregulation of type 3 inositol (1, 4, 5)-trisphosphate receptor decreases breast cancer cell migration through an oscillatory Ca2+ signal”. In: Oncotarget 8.42, p. 72324.

Noble, Denis (1960). “Cardiac action and pacemaker potentials based on the Hodgkin-Huxley equations”. In: Nature 188.4749, pp. 495–497.

Oeggerli, Martin et al. (2012). “Role of KCNMA1 in breast cancer”. In: PloS one 7.8, e41664.

Onkal, Rustem et al. (2008). “Alternative splicing of Nav1. 5: an electrophysiological comparison of ‘neonatal’and ‘adult’isoforms and critical involvement of a lysine residue”. In: Journal of cellular physiology 216.3, pp. 716–726.

Panner, Amith and Robert D Wurster (2006). “T-type calcium channels and tumor proliferation”. In: Cell calcium 40.2, pp. 253–259.

Pieri, Massimo et al. (2013). “Over-expression of N-type calcium channels in cortical neurons from a mouse model of Amyotrophic Lateral Sclerosis”. In: Experimental neurology 247, pp. 349–358.

Prevarskaya, Natalia, Roman Skryma, and Yaroslav Shuba (2018). “Ion channels in cancer: are cancer hallmarks oncochannelopathies?” In: Physiological reviews 98.2, pp. 559–621.

Quicke, Peter et al. (2022). “Voltage imaging reveals the dynamic electrical signatures of human breast cancer cells”. In: Communications Biology 5.1, p. 1178.

Rehak, Renata et al. (2013). “Low voltage activation of KCa1. 1 current by Cav3-KCa1. 1 complexes”. In: PloS one 8.4, e61844.

Ribeiro, Mafalda et al. (2020). “Human breast cancer cells demonstrate electrical excitability”. In: Frontiers in neuroscience 14, p. 404.

Rizaner, Nahit et al. (2016). “Intracellular calcium oscillations in strongly metastatic human breast and prostate cancer cells: control by voltage-gated sodium channel activity”. In: European Biophysics Journal 45, pp. 735–748.

Roger, Sébastien, Pierre Besson, and Jean-Yves Le Guennec (2003). “Involvement of a novel fast inward sodium current in the invasion capacity of a breast cancer cell line”. In: Biochimica et Biophysica Acta (BBA)-Biomembranes 1616.2, pp. 107–111.

Roger, Sébastien, Marie Potier, et al. (2004). “Description and role in proliferation of iberiotoxin-sensitive currents in different human mammary epithelial normal and cancerous cells”. In: Biochimica et Biophysica Acta (BBA)-Biomembranes 1667.2, pp. 190–199.

Rüdiger, S, JW Shuai, and IM Sokolov (2010). “Law of mass action, detailed balance, and the modeling of calcium puffs”. In: Physical review letters 105.4, p. 048103.

Selimi, Zoja et al. (2023). “A detailed analysis of single-channel Nav1. 5 recordings does not reveal any cooperative gating”. In: The Journal of Physiology 601.17, pp. 3847–3868.

Sizemore, Gina et al. (2020). “Opening large-conductance potassium channels selectively induced cell death of triple-negative breast cancer”. In: BMC cancer 20, pp. 1–14.

Smith, Greg Conradi (2019). Cellular biophysics and modeling: a primer on the computational biology of excitable cells. Cambridge University Press.

Strogatz, Steven H (2018). Nonlinear dynamics and chaos with student solutions manual: With applications to physics, biology, chemistry, and engineering. CRC press.

Szatkowski, Cécilia et al. (2010). “Inositol 1, 4, 5-trisphosphate-induced Ca 2+ signalling is involved in estradiol-induced breast cancer epithelial cell growth”. In: Molecular cancer 9, pp. 1–13.

Taylor, James T et al. (2008). “Selective blockade of T-type Ca2+ channels suppresses human breast cancer cell proliferation”. In: Cancer letters 267.1, pp. 116–124.

Tsaneva-Atanasova, Krasimira et al. (2007). “Mechanism of spontaneous and receptor-controlled electrical activity in pituitary somatotrophs: experiments and theory”. In: Journal of Neurophysiology 98.1, pp. 131–144.

Winkler, Frank et al. (2023). “Cancer neuroscience: State of the field, emerging directions”. In: Cell 186.8, pp. 1689–1707.

Wolf, Anne and Gunther Wennemuth (2014). “Ca2+ clearance mechanisms in cancer cell lines and stromal cells of the prostate”. In: The Prostate 74.1, pp. 29–40.

Yamada, Walter M, Christof Koch, and Paul R Adams (1989). “Multiple channels and calcium dynamics”. In: Methods in neuronal modeling 1989, pp. 137–170.

Yang, Ming et al. (2020). “Voltage-dependent activation of Rac1 by Nav1. 5 channels promotes cell migration”. In: Journal of cellular physiology 235.4, pp. 3950–3972.

Zhang, Zheng et al. (2013). “Kinetic model of Nav1. 5 channel provides a subtle insight into slow inactivation associated excitability in cardiac cells”. In: PloS one 8.5, e64286.

